# Detailed single-cell mapping of the transcriptional response to a virus infection driven by copy-back viral genomes

**DOI:** 10.64898/2026.01.06.697933

**Authors:** Yanling Yang, Emna Achouri, Munyaradzi Tambo, Carolina B. López

## Abstract

The antiviral response to several clinically significant viruses, including respiratory syncytial virus and parainfluenza virus, is driven by copy-back viral genomes (cbVGs) generated during virus replication. However, the broader impact of cbVGs on the functional states of host cells remains undefined. Here, we developed a single-cell RNA-sequencing and computational framework to map cbVG-driven host responses during Sendai virus infection. Unsupervised profiling identified distinct transcriptional states throughout the course of infection, highlighting a shift from early antiviral signaling to later inflammatory and remodeling programs. Stratifying infected cells by cbVG status demonstrated that cbVG-positive cells initiate interferon and chemokine programs, which later spread to cbVG-negative cells. At later stages, cbVG-positive cells acquire additional signaling, cytoskeletal, transcriptional, and stress-adaptation programs, which are absent in cbVG-clean infection. This work defines the broader cbVG-driven layered and dynamic host response and provides a valuable high-resolution resource of the temporal cellular response to a virus infection.

## Introduction

Non-standard defective viral genomes (nsVGs) are generated in nearly all RNA virus infections and have long been recognized as potent activators of innate immunity^1–3^. Among them, copy-back viral genomes (cbVGs) emerge during negative-sense RNA virus replication and strongly induce the antiviral interferon response through triggering signaling by cytosolic RNA sensors^4–12^. Although cbVGs have been detected across multiple virus families and shown to impact infection dynamics and clinical outcomes^7,13–25^, how cbVGs shape host responses through the infection course remains unresolved. A fundamental barrier has been the lack of single cell data that allows building a conceptual model to explain how a rare cbVG-producing subpopulation of infected cells drives antiviral programs across an infected tissue. So far, studies on the impact of cbVGs in antiviral immunity have either relied on bulk measurements of infected populations^26–30^, which obscure cell-to-cell heterogeneity and mask cell-state transitions, or have focused mostly on the expression of interferons^31,32^ overlooking the broader impact of cbVG-driven signals through time and across the infected population.

The biology of cbVGs has primarily been investigated after preparing cbVG-high virus stocks or adding purified cbVGs to infected cultured cells^33,34^, or after isolating cbVG-rich populations by RNA-FISH and flow cytometry^35^. While these approaches established cbVGs as strong inducers of antiviral signaling, they lack the resolution to distinguish cbVG-positive (cbVG⁺) cells from those responding indirectly (cbVG⁻) within the same population. Moreover, cbVG formation is stochastic producing diverse species that vary in abundance across cells^14^. As a result, early signaling events initiated by a small cbVG⁺ subpopulation are diluted in bulk measurements, preventing analysis of temporal dynamics, paracrine effects, and cbVG-driven programs that emerge later in infection. Together, these limitations have left several gaps in the understanding of the early events of antiviral immunity, including understanding how cbVGs orchestrate population-wide responses, dissecting transcriptional programs directly driven by cbVGs versus secondary effects of antiviral signaling, and importantly, understanding the full impact of cbVGs on the infected cell.

Recent advances in single-cell RNA sequencing (scRNA-seq) enable simultaneous profiling of host and viral transcriptomes in thousands of individual cells^36–40^. Single-cell approaches have revealed heterogeneity in viral gene expression, innate immune activation, and cell-state transitions during infection^41–44^. However, direct detection of cbVGs at single-cell resolution remains technically challenging. CbVGs can only be distinguished from the standard viral genome by recognizing the junction sequence that forms when the viral polymerase falls off the template strand during replication and rejoins the nascent RNA strand to continuing copying this strand of the genome. In addition, most RNA viruses produce populations of cbVG that differ in junction positions^14,17,18,22,45–47^. Standard scRNA-seq relies on identification of polyadenylated transcripts and captures only one end of transcripts near the template-switch oligo^48,49^. As cbVGs are not polyadenylated, and cbVG junction sequences vary in their position relative to the template-switch oligo, most cbVGs are lost during standard scRNA-seq library preparation.

Here, we took advantage of the murine parainfluenza virus Sendai strain Cantell (SeV Cantell) to develop an integrated single-cell sequencing and computational framework to directly identify cbVGs in individual cells during infection. SeV C produces a single dominant cbVG (cbVG-546) that potently induce interferon expression^13^. Importantly the junction sequence of cbVG-546 is detectable by the conventional 10x Genomics scRNA-Seq approach. Unsupervised profiling uncovered dynamic cellular states of SeV-positive cells, revealing cbVG-associated transitions from early interferon responses to later inflammatory programs. By resolving SeV-positive cells into cbVG⁺, cbVG⁻ across time, we define cbVG-driven transcriptional programs. Single-cell profiling at 6, 12, and 24 hours post infection revealed that cbVGs initiate interferon and chemokine programs from a rare cbVG⁺ subpopulation at early time points, which subsequently propagate to cbVG⁻ cells. At later time points, cbVG⁺ cells express additional transcriptional modules that remain poorly represented in cbVG⁻ cells, defining cbVG-specific programs beyond interferon responses. These results revealed a layered host response to virus infection driven by cbVGs, provide a single-cell roadmap for studying the impact of cbVGs in other infections and serve as a resource for dissecting the components of the antiviral response within the first 24 h of infection.

## Results

### Single-cell sequencing enables direct detection of cbVGs in individual infected cells

cbVGs are generated by the viral RNA-dependent RNA polymerase during genome replication and contain a unique rejoin junction sequence that distinguishes them from standard viral genomes (Fig. S1). The VODKA2 pipeline detects cbVGs from short read RNA-seq datasets by identifying these junction-spanning reads (Fig. S1B). To directly resolve cbVG-producing cells in single-cell data, we modified the standard scRNA-seq workflow by incorporating a virus-specific reverse transcription primer^50–52^ in addition to the standard oligo(dT) priming for capturing mRNA (Fig. 1A). The virus-specific primer was designed to capture viral transcripts including the junction sequence of cbVG-546, the dominant cbVG species generated during infection with SeV C^13^. Both primers share a common 5’ overhang sequence, enabling recovery of host mRNAs, viral mRNAs, antigenomes, and cbVG transcripts during library preparation (Fig. 1B). Junction-spanning reads from cbVG-546 can be captured because the transcript containing the junction is 278 bp in length and the rejoin site is positioned 94 bp from the template-switch oligo, within the read length of 10x-Illumina sequencing (Fig. 1C). To characterize cbVG-driven responses over time, A549 cells were infected with SeV C at a multiplicity of infection (MOI) of 1.5-2 and sampled at 6, 12, and 24 hpi. An independent infection at higher MOI (MOI=5) was performed to increase the probability of detecting early cbVG-induced programs and to test reproducibility across infection settings. Sequencing data were processed by integrating a standard scRNA-seq pipeline with VODKA2 (Fig. S2). The number of cells passing quality control, as well as the number of SeV⁺ and cbVG⁺ cells per condition, are shown in Fig. 1D. Although the number of infected cells was similar across conditions, the number of cbVG⁺ cells increased with time and MOI, consistent with progressive cbVG accumulation during infection. Viral UMI counts increased with MOI at 6 hpi, and with infection time at later time points, with 1-30 cbVG UMI per cbVG⁺ cell detected across samples (Fig. 1E). Together, these results demonstrate that our modified scRNA-seq approach reliably detects viral and cbVG reads across infection time points at single-cell resolution, enabling direct identification of cbVG⁺ cells during SeV infection.

**Fig. 1.**
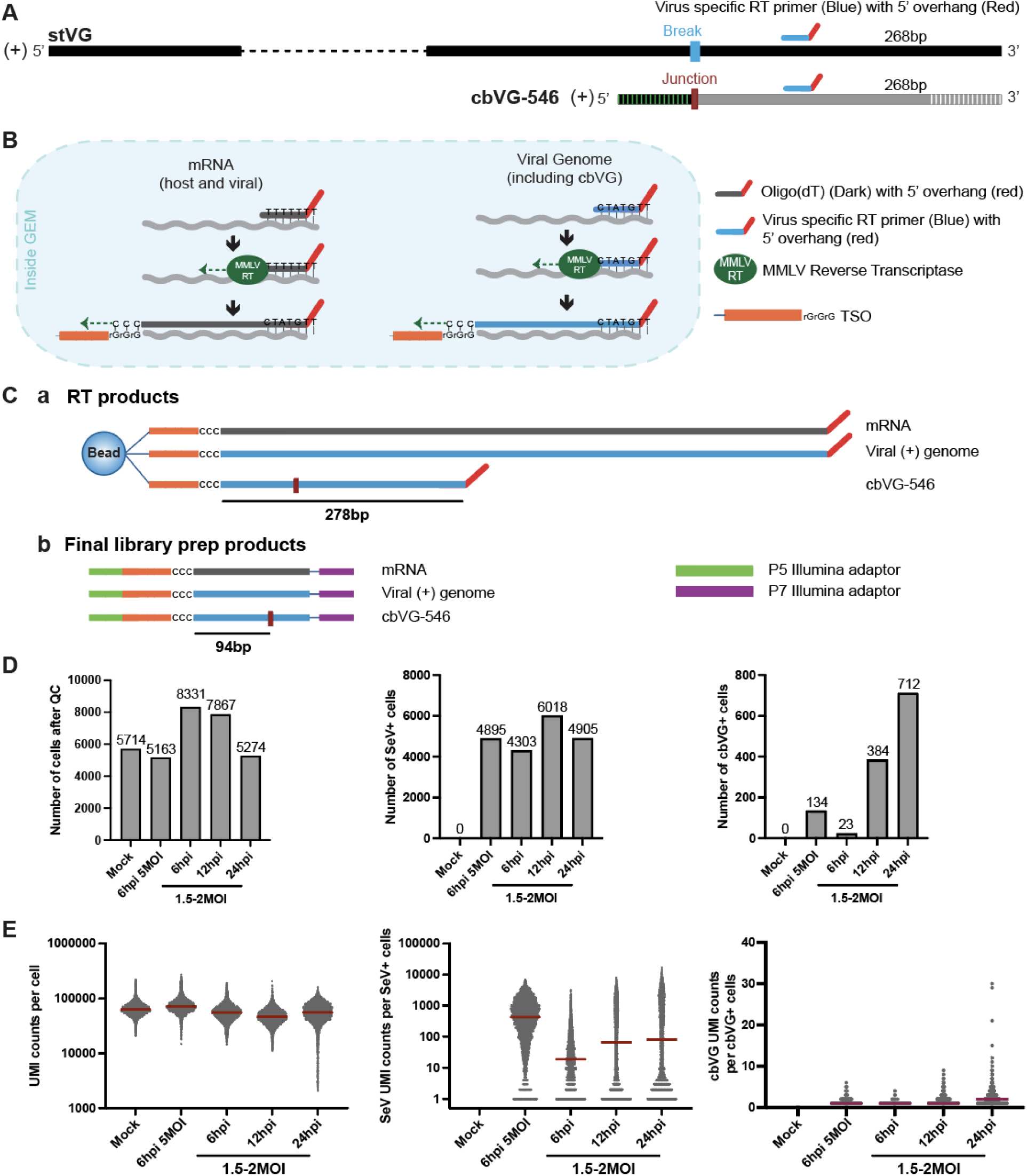
Detection of cbVGs at the single-cell level. (A) A virus-specific RT primer (blue) with a 5′ overhang (red) was designed to bind 256 bp from the 3′ end of the viral genome to capture both stVGs and cbVG-546. (B) Diagram showing mRNA and viral RNA capture and reverse transcription (RT) inside 10x Genomics Chromium droplets. (C) Illustration of RT and library preparation for Illumina sequencing. (a) RT products of mRNA, viral genome, and cbVG-546 with the cbVG junction region labeled. A 278 bp of cbVG-546 that includes the junction region is transcribed. (b) Final library prep products with Illumina adaptors (green and purple) added to both ends. The cbVG-546 junction is 94 bp from the TSO. (D) Number of cells passing quality control (QC) and numbers of SeV-positive and cbVG-positive cells in each sample. (E) Total UMI counts, SeV reads, and cbVG reads per cell in each sample.

### Single-cell analysis identifies a rare cbVG⁺ subpopulation as the origin of early interferon induction

To define the relationship between the different types of viral genomes and early interferon responses, we performed unbiased clustering of infected cells based on their transcriptomes and overlaid viral RNA content, including both standard viral genomes and cbVG junction reads (Fig. 2). While SeV RNA was distributed across all clusters, cbVGs were enriched in a subset of transcriptional states, indicating that cbVG production is not uniform across infected cells (Fig. 2A-B). A distinct IFN-producing cluster was identified under all conditions, characterized by expression of IFNB1 and IFNL1-3 among the top marker genes (Fig. 2C-D). Although this cluster contained a high proportion of SeV⁺ cells, other clusters with similarly high infection rate did not express IFNs, indicating that infection alone is insufficient to initiate interferon production (Fig. 2D-E) and confirming published reports of a the lack of correlation between viral load and IFN expression^6,7,21,53^.

**Fig. 2.**
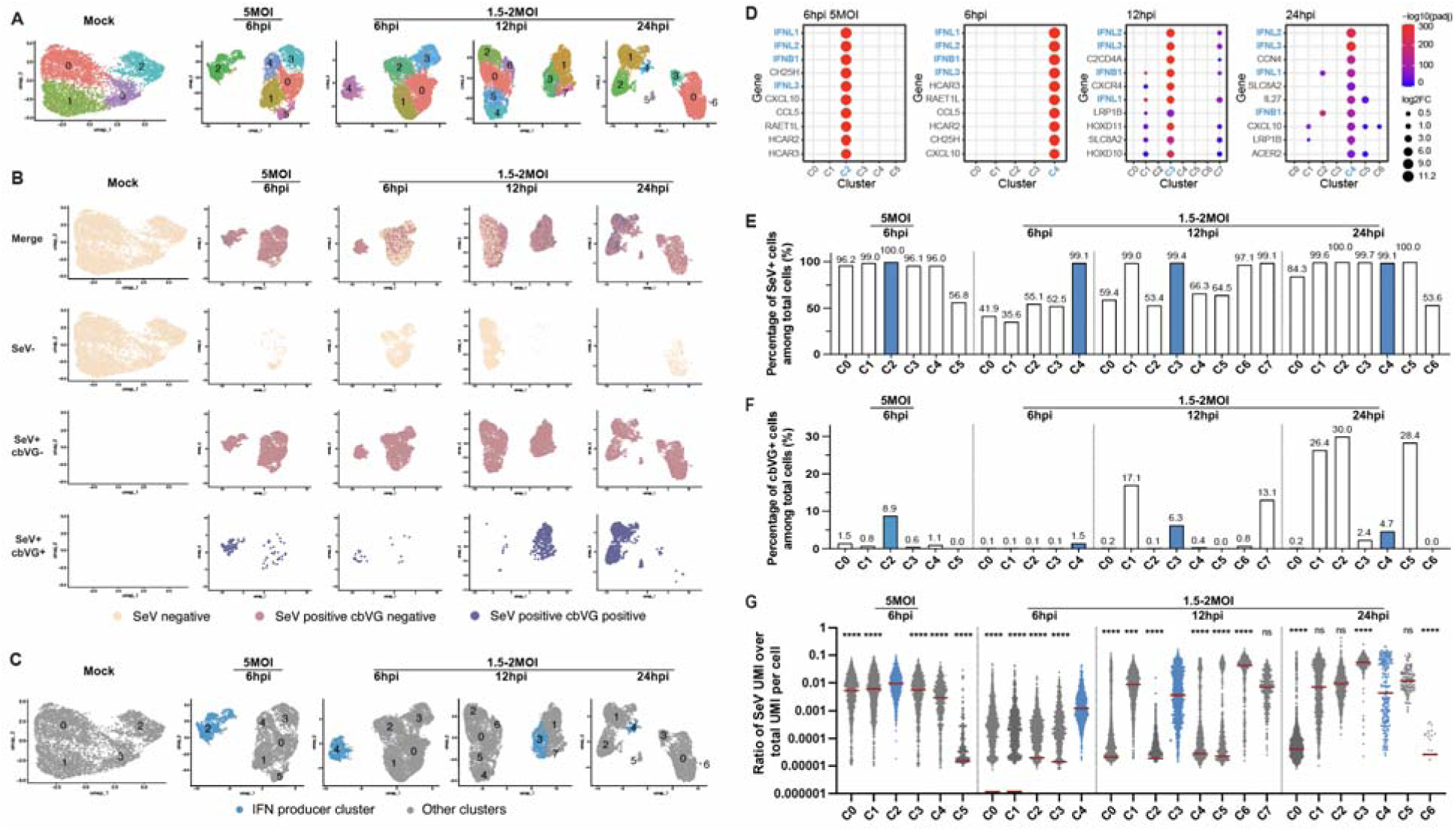
Unbiased clustering of each sample and identification of IFN-producing clusters in SeV-infected cells. (A) UMAPs showing unbiased clustering of mock and SeV-infected A549 cells at different infection conditions (5 MOI or 1.5-2 MOI) and time points (6, 12, and 24 hpi). Each color represents a distinct cell cluster in each sample. Clusters in different samples are not comparable based on their color. (B) UMAPs of SeV^-^, SeV^+^cbVG^-^, and SeV^+^cbVG⁺ cells in each sample. Cells are colored according to viral RNA detection status: SeV^-^ (beige), SeV^+^cbVG⁻ (pink), and SeV^+^cbVG⁺ (slate blue). (C) UMAPs showing IFN-producing (blue) clusters across mock and SeV-infected samples under different infection conditions (5 MOI or 1.5-2 MOI) and time points (6, 12, and 24 hpi). (D) Expression levels of the top 10 expressed marker genes of the IFN-producing clusters in different infection conditions. (E-F) Percentages of SeV^+^ cells (E) and cbVG⁺ cells (F) among total cells in each cluster. (G) Abundance of SeV reads in each cluster represented as the ratio of SeV UMI counts over total UMI counts per cell. Each dot represents a single cell. Statistical differences in the median values across all clusters in each sample were analyzed using the Kruskal-Wallis test. Following a significant overall test, pairwise comparisons were performed using Dunn’s post-hoc test, in which each cluster was compared specifically against the IFN-producing cluster. Significance levels are indicated as ns for not significant and asterisks for significant differences (***p < 0.001 and ****p < 0.0001).

SeV cbVG⁺ cells were consistently present within the IFN-producing cluster across MOIs and time points. At 6 hpi, the IFN-expressing cluster showed a higher average viral load than other clusters and higher counts of cbVGs (Fig. 2D-G) consistent with the requirement of viral replication machinery for cbVG accumulation^3,24,54^. This association diminished at later timepoints, likely explained by cbVG-driven interference with viral replication^3,55^. By 12 hpi, some cbVG⁺ clusters lacked IFN expression (Fig. 2E-G) likely reflecting host-mediated shutdown of interferon signaling and/or the transition of infected cells into different transcriptional states. Notably, many SeV+ cells across clusters had similar or higher total viral RNA abundance than cells in the IFN cluster, indicating that cbVG detection does not simply reflect viral load and demonstrating that cbVGs, rather than viral genome abundance, are the strongest correlate with interferon expression during early infection. Together, these results support data from bulk cell populations showing that cbVGs are the primary triggers of early antiviral programs^4,7–9,13,47,56,57^ and establish that cbVGs are the initial drivers of antiviral signaling rather than it being a correlate of viral genome abundance.

### Distinct dynamic cellular transcriptional states are identified during SeV infection

To resolve cellular transcriptional profiles reflecting distinct cellular states across infection conditions, we performed integrated analysis followed by unsupervised clustering of SeV⁺ cells. This analysis revealed five distinct transcriptional profiles (C0-C4) that captured the dominant cellular programs present during the first 24 h of infection (Fig. 3A). Each state was characterized by distinct gene expression programs (Fig. 3B-D), representing metabolic adaptation, cell-cycle regulation, epithelial secretory responses, interferon production, and inflammatory signaling. C0 expressed genes associated with metabolic and oxidative stress-response programs (e.g., HMOX1, SLC40A1, ACOX2), consistent with iron homeostasis, lipid metabolism, and redox balance (Fig. S3A). C1 was enriched for cell-cycle and chromatin organization programs (e.g., HIST1H1B, CENPA, CCNA2), reflecting DNA replication and mitotic progression (Fig. S3B). C2 was enriched in epithelial adhesion and secretory machinery genes (e.g., CEACAM6, MUC5AC, CLDN2, LCN2) together with cytokine-mediated signaling and immune activation programs (Fig. S3C). C3 represented the interferon-producing state, defined by high expression of IFNB1 and IFNL1-3, and enriched for antiviral defense programs, including type I/III interferon signaling (Fig. S3D). C4 exhibited inflammatory signaling modules, including chemokine induction and was enriched in NF-κB-associated signaling (e.g., CCL17, ICAM1, ACKR1) (Fig. S3E). We then analyzed the expression levels of cluster-defining marker genes relative to mock and assessed the proportion of cells expressing them in each cluster at each experimental condition (Fig. 3C-D). All clusters contained more than 3,000 cells, enabling reliable assessment of gene expression levels and the proportion of marker-positive cells across conditions (Fig. S4A). We observed that expression of metabolic and oxidative stress-related genes such as HMOX1, ACOX2, and CYP24A1, as well as cell-cycle and chromatin organization genes including HIST1H1B, CENPA, and CCNA2, were significantly downregulated in the IFN-producing (C3) and inflammatory (C4) clusters. In contrast, IFNL1, IFNL2, IFNL3, and IFNB1 were strongly upregulated in both C3 and C4).

**Fig. 3.**
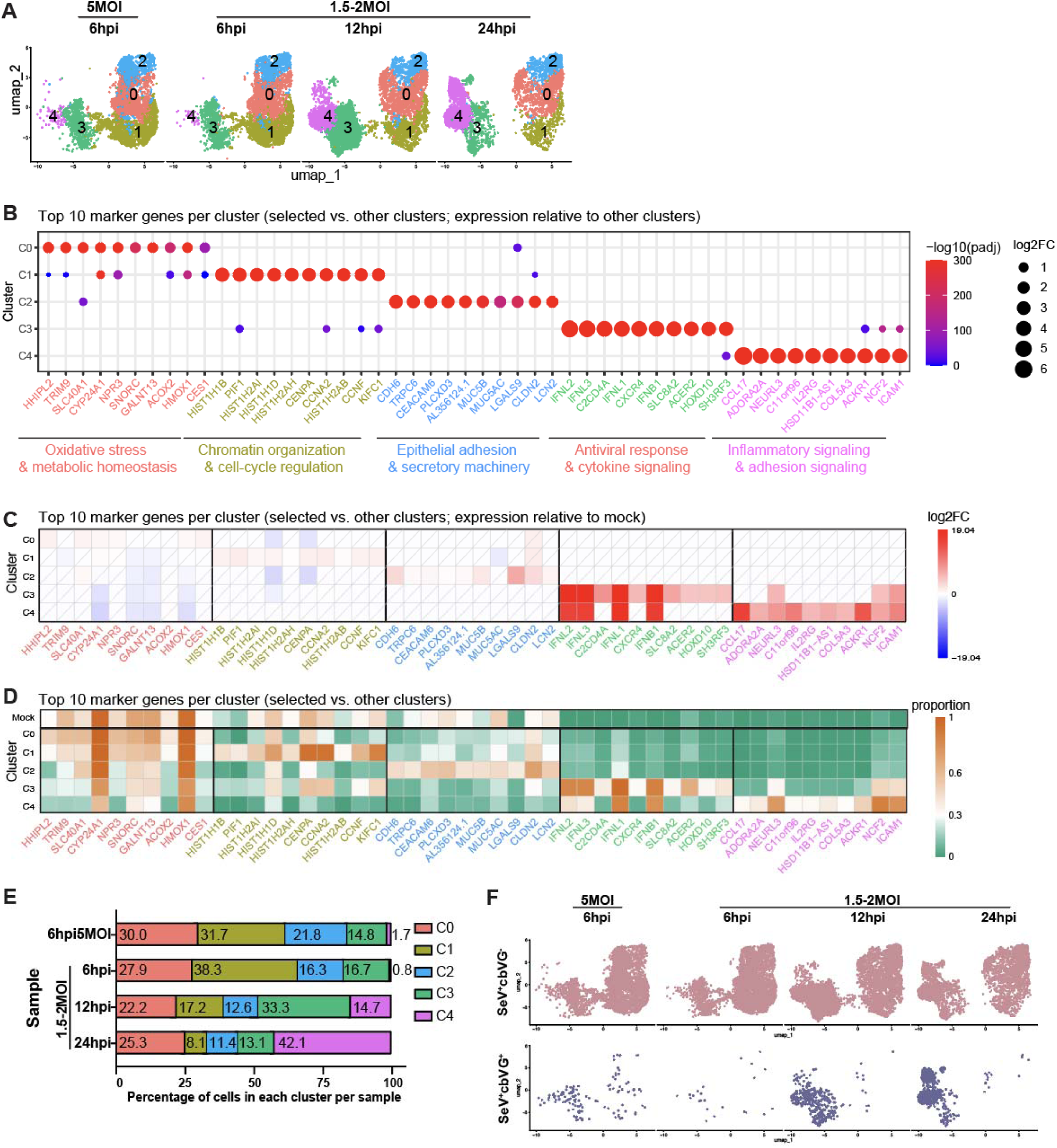
SeV+ cells can be divided into five populations during the first 24 h f infection based on transcriptome signatures. (A) Integrated analysis of all SeV+ cells from each sample. (B) Bubble plot showing the top 10 marker genes for each SeV+ cluster (C0-C4) identified relative to other clusters. Dot size represents log2 fold change, and color indicates statistical significance (-log10 adjusted p value). Functional categories of marker genes are labeled below. (C) Heatmap showing the expression of the same top 10 marker genes in each cluster relative to mock-infected cells, with the color scale indicating log2 fold change. White tiles with diagonal lines indicate features that did not meet the predefined adjusted P-value and proportion thresholds. (D) Heatmap showing the proportion of cells expressing each marker gene across clusters and mock (uninfected sample), with the color scale representing expression proportion. (E) Percentage of cells in each population across samples. (F) UMAPs showing the distribution of SeV^+^ cells under different infection conditions (5 MOI or 1.5-2 MOI) and time points (6, 12, and 24 hpi). Separate panels show SeV^+^cbVG^−^ (pink) and SeV^+^cbVG⁺ (slate blue) populations individually.

We next quantified the distribution of cells across states over time (Fig. 3E). While the proportion of C0 remained relatively constant, the other four transcriptional profiles displayed marked temporal dynamics. C1 was most abundant at 6 hpi and decreased thereafter, indicating suppression of proliferative programs as infection progressed. C2 diminished over time indicating reduction of cells with active adhesion and secretory pathways. C3 (interferon-producing) peaked at 12 hpi, consistent with early antiviral activation. By contrast, C4 expanded dramatically from ∼1% at 6 hpi to 42% at 24 hpi, indicating a population shift toward late inflammatory programs. Analysis of SeV⁻ cells did not reveal comparable dynamics, as the top 20 marker genes in each cluster were predominantly interferon-stimulated genes (ISGs) and were upregulated across clusters relative to mock (Fig. S5), demonstrating that these transitions are specific to infected cells. To relate these transcriptional profiles to the presence of cbVGs, we mapped cbVG⁺ cells across clusters (Fig. 3E and S4B). cbVG⁺ cells were concentrated in C3 and C4, with C3 containing most cbVG⁺ cells early in infection and C4 becoming the dominant cbVG⁺ population at 24 hpi. This pattern aligns with the temporal progression from early interferon induction (C3) to a later broader inflammatory response (C4).

Together, these analyses identify five functional transcriptional programs within a clonal population of infected cells, four of which change quantitatively within the first 24 h of infection. cbVG⁺ cells show primarily antiviral and inflammatory states, and their redistribution over time highlights a trajectory from early interferon production to late inflammatory signaling. Notably, metabolic and cell-cycle programs (C0 and C1), as well as cell adhesion and secretory programs (C2) were suppressed in C3 and C4 when comparing with mock, indicating that activation of antiviral programs coincides with downregulation of metabolic, proliferative and other processes, consistent with a cell-state transition toward a stress-adapted antiviral phenotype.

### cbVGs impact the dynamics of the antiviral response within a population of infected cells

To more directly evaluate how cbVGs shape host responses within infected populations, we stratified SeV⁺ cells from each sample into cbVG⁺ and cbVG⁻ subgroups. The number of cells in each group and the SeV UMI counts per cells are shown in Fig. S6. Then we compared transcriptional changes in cbVG⁺ and cbVG⁻ cells relative to mock controls (Supplementary Table 1-2, and Fig. S7). cbVG⁺ cells displayed a greater number of differentially expressed genes at all time points and conditions than cbVG⁻ cells, except at the earliest stage of infection at low MOI (Fig. 4A-B, indicating an overall stronger transcriptional response associated with cbVG presence. When combining the top upregulated genes from all groups (Fig. 4C-D, interferons and chemokines (e.g., IFNB1, IFNL1-3, CCL5, CXCL10) were exclusively found in cbVG⁺ cells during early infection and became broadly expressed in both cbVG⁺ and cbVG⁻ cells at later time points. This observation is consistent with IFN priming and sensitization of bystander cells to response to the infection. In contrast, a subset of genes, including CCL3 and GNG8, were induced only in cbVG⁺ cells at later timepoints, suggesting a temporal split on cbVG-driven transcriptional responses. Classic interferon-stimulated genes (ISGs), such as OASL, IFIT1, and ISG15, were induced in both cbVG⁺ and cbVG⁻ cells from 6 hpi onward, reflecting autocrine and paracrine responses to IFNs (Fig. 4C-D). A larger proportion of downregulated genes was observed at 12 and 24 hpi relative to 6 hpi (Fig. 4E-F), consistent with broad suppression of metabolic and proliferative programs as infection progressed.

**Fig. 4.**
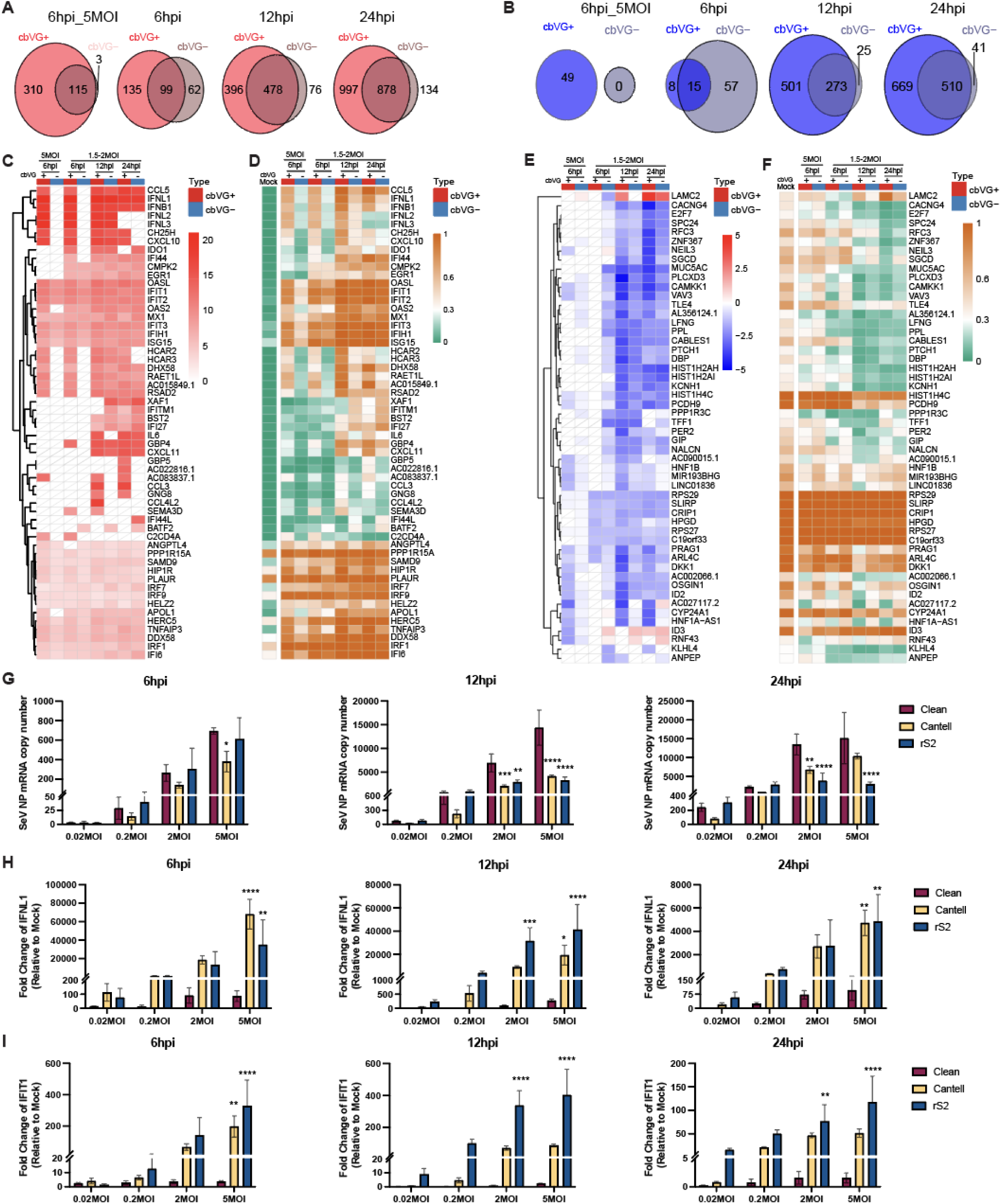
Viral RNA levels and cbVG-associated gene expression across infection conditions. (A-B) Venn diagrams showing the overlap of DEGs between cbVG⁺ and cbVG⁻ cells compared with mock-infected cells at 6hpi (5MOI), 6hpi, 12hpi, and 24hpi. Numbers indicate unique and shared genes between the two groups. (A) Upregulated genes (pct.cbVG⁺ > 0.3, padj < 0.01, and log2FC > 1). (B) Downregulated genes (pct.mock > 0.3, padj < 0.01, and log2FC < -1). (C-F) Heatmaps showing the expression levels (C and E) and the proportion of expressing cells (D and F) for the top 20 genes significantly upregulated (C-D) or downregulated (E-F) in cbVG⁺ or cbVG⁻ cell populations in each sample. The color scale indicates the average log2 fold change (C and E) or the proportion of cells expressing each gene (D and F). White tiles with diagonal lines indicate features that did not meet the predefined adjusted P-value and proportion thresholds. (G) SeV NP, (H) IFNL1, and (I) IFIT1 expression in A549 cells infected with SeV cbVG clean stock or cbVG-high stocks (Cantell and rS2) at indicated MOIs (0.02, 0.2, 2, and 5) and time points (6, 12, and 24 hpi). Expression of mRNA was calculated relative to the housekeeping index with GAPDH and β-actin. IFNL1 and IFIT1 expression represents fold change relative to mock-infected controls. Data represent means ± SD of three biological replicates. Data were analyzed using two-way ANOVA, followed by Šídák’s multiple-comparisons test to compare Clean versus Cantell or rS2. Adjusted P values are shown. *, P < 0.05; **, P < 0.01; ***, P < 0.001; ****, P < 0.0001.

To validate whether these genes were induced by cbVGs rather than by SeV infection alone, we quantified using qPCR SeV NP, IFNL1, and IFIT1 expression in cells infected with a cbVG-clean stock prepared by rescuing the virus by reverse genetics, a recently described cbVG-high stock rS2 with different composition of cbVGs^14^, and SeV Cantell that contains cbVG546 (Fig. 4G-I). NP levels remained high in cells infected with the clean stock across time points and MOIs, consistent with productive viral replication in the absence of pre-existing cbVGs (Fig.4G). In contrast, IFNL1 and IFIT1 were not upregulated in clean stock infections but showed robust induction in cells infected with different cbVG-high stocks at multiple time points (Fig.4H-I), confirming that their activation is driven by cbVG presence rather than the viral infection itself. These results provide orthogonal validation of the single-cell findings, confirming that key antiviral genes are selectively induced in the presence of cbVGs. Together with the transcriptomic data, they show that cbVGs determine both the timing and amplitude of early antiviral induction by triggering interferon and chemokine programs in a rare cbVG⁺ subpopulation, which then propagate to cbVG⁻ cells through paracrine signaling. Beyond this shared antiviral phase, cbVG⁺ cells sustain a distinct subset of late transcriptional programs, demonstrating that cbVGs also drive divergent cell-state outcomes later in infection.

### cbVG induce distinct early and late transcriptional programs in infected cells

To more comprehensively define the transcriptional programs driven by cbVG, we then identified all genes that were significantly upregulated over mock in cbVG⁺ cells, but not cbVG⁻ cells, at different times post infection. We identified 43 genes upregulated at 6 hpi regardless of MOI (Fig. 5). The genes in this category included interferons (IFNB1, IFNL1-3) and the chemokines CCL5 and CXCL10, which were strongly induced exclusively in cbVG⁺ cells (Fig. 5A). In contrast, in cbVG⁻ cells these genes were not upregulated, despite cel cells being infected. However, by 12 hpi, most of these genes were expressed in both cbVG⁺ and cbVG⁻ cells, and remained elevated at 24 hpi, indicating that the broader infected population express these antiviral effectors at later time points (Fig. 5A). This temporal shift from cbVG-restricted to population-wide expression is consistent with enhanced responsiveness of cells to the infection, wherein IFNs and chemokines produced by a small cbVG⁺ subset signal surrounding cells to be primed to respond to the infection. We classified these early induced genes into seven functional programs based on known or predicted functions: interferon responses; chemokine signaling; metabolic and lipid signaling; complement and innate immune amplification; cell–cell interaction; transcriptional and stress-response regulation; and uncharacterized genes with predicted immune roles. Of note, because for this analyses we only considered genes expressed at both MOIs tested for a more robust analysis, we left out several genes that are sdetectable only at high MOI, including OAS2, TRAF1, and PAK3 (Fig. S8). Overall, these data show that early cbVG⁺ cells induce multifunctional antiviral responses rich in immune communication signals.

**Fig. 5.**
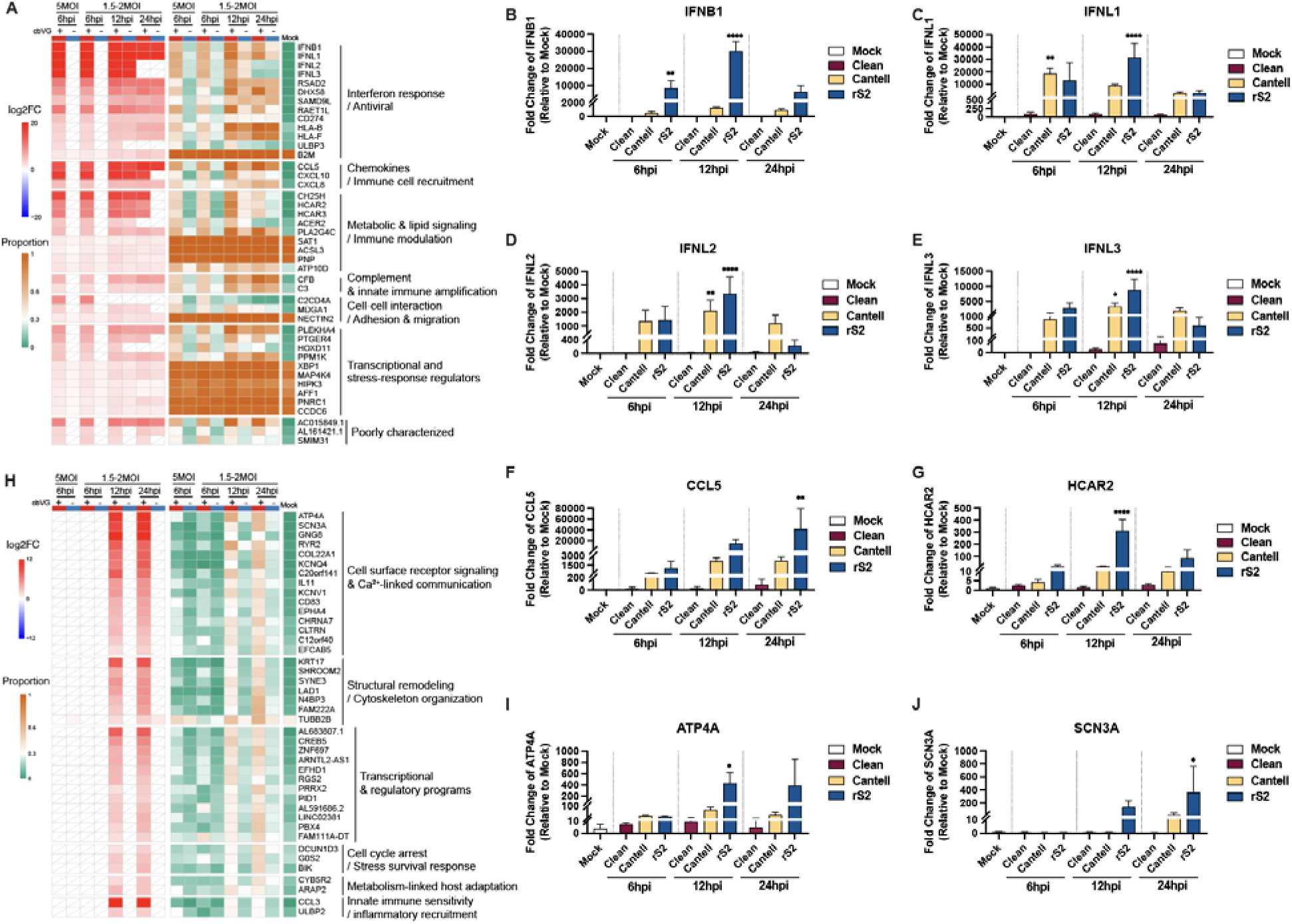
Functional classification and validation of genes upregulated exclusively in cbVG cells. (A) Heatmap showing genes specifically upregulated in cbVG⁺ but not cbVG⁻ cells at both 6hpi samples. Genes were grouped into seven functional categories based on their known or predicted biological roles. The red-blue color scale indicates the log2 fold change (log2FC), while the orange-green color scale represents the proportion of cells expressing each gene. White tiles with diagonal lines indicate features that did not meet the predefined adjusted P-value and proportion thresholds. (B-G) Representative genes validated by qPCR. A549 cells were infected with SeV Cantell stocks containing different cbVG compositions: cbVG clean stock, Cantell: high level of cbVG-546, rS2: high levels of cbVGs other than cbVG-546. Each gene mRNA levels were quantified by qPCR and are presented as fold change relative to mock-infected samples. Cells collected at 6, 12, and 24 hpi. Ordinary one-way ANOVA followed by Dunnett’s multiple comparisons test comparing each infection group with the mock control (n = 3). Adjusted P values are shown. Significance P values are indicated as follows: P < 0.05 (*), < 0.01 (**), < 0.001 (***), < 0.0001 (****). (H) Heatmap showing genes specifically upregulated in cbVG⁺ but not cbVG− cells at both 12hpi and 24hpi samples. Genes were grouped into six functional categories based on their known or predicted biological functions. The red-blue color scale indicates the log2 fold change (log2FC), while the orange-green color scale represents the proportion of cells expressing each gene. White tiles with diagonal lines indicate features that did not meet the predefined adjusted P-value and proportion thresholds. (I-J) Representative genes validated by qPCR. A549 cells were infected with SeV Cantell stocks containing different cbVG compositions: cbVG clean stock, Cantell, and rS2. Each gene mRNA levels were quantified by qPCR and are presented as fold change relative to mock-infected samples. Cells were collected at 6, 12, and 24 hpi. Data represent mean ± SD from three biological replicates. Ordinary one-way ANOVA followed by Dunnett’s multiple comparisons test comparing each infection group with the mock control (n = 3). Adjusted P values are shown. Significance P values are indicated as follows: P < 0.05 (*), < 0.01 (**), < 0.001 (***), < 0.0001 (****).

To validate these data, we next performed infections with a cbVG-clean stock, and two cbVG-high stocks, Cantell and rS2, and tested for gene expression by qPCR. Our data confirmed that early induced genes (Fig. 5B-G, S9A-C) were detectable only in cbVG-high infections and were absent in infections with the cbVG-clean stock. These data mirror the scRNA-seq results and demonstrate that early antiviral gene induction is cbVG-dependent and is not specific to the cbVG-546 species and can be induced by diverse cbVG populations. Together, these findings show that early antiviral signaling originates from a rare population of cbVG⁺ cells, which acts as the initial source of interferon and chemokine production and initiate a population-wide antiviral response through paracrine amplification.

To characterize the later cbVG-driven transcriptional programs, we analyzed genes that were upregulated only in cbVG⁺ cells at 12 and 24 hpi, after excluding those already induced at 6 hpi (Fig. 5H). This analysis revealed 105 genes that define a late cbVG-dependent program. Among the top 41 genes (log_2_FC > 2) were ATP4A, SCN3A, KRT17, and CCL3, which were highly and specifically enriched in cbVG⁺ cells at late time points. These late-phase cbVG⁺ programs encompass cell-surface receptor signaling, Ca²⁺-linked communication, cytoskeletal remodeling, transcriptional regulation, stress-survival mechanisms, and innate immune sensitivity, indicating that cbVG⁺ cells undergo distinct cell-state transitions that are not shared by the broader population in the time frame analyzed. Genes that were transiently upregulated at 12 hpi or 24 hpi are shown in Fig. S10. qPCR validation confirmed that these late-induced genes were not upregulated at 6 hpi but became strongly induced in infections with SeV Cantell and rS2 at later time points. These transcripts remained absent in infections with cbVG-clean stock (Fig. 5I-J, S9D-F. These results indicate that cbVG⁺ cells do not only initiate antiviral programs but later acquire a second wave of transcriptional responses that reflect cell-state remodeling beyond interferon defenses.

### Several antiviral programs are shared across the infected cell population

We next defined the shared antiviral response by identifying genes that were upregulated in both cbVG⁺ and cbVG⁻ cells throughout the infection course (Fig. 6). We identified 70 genes that were significantly upregulated in both 6 hpi samples (Fig. 6A), and nearly all remained elevated during 12 and 24 hpi, revealing a stable shared antiviral program. These broadly induced genes clustered into five expression programs based on temporal and functional features: (1) canonical ISGs; (2) interferon-responsive non-canonical genes; (3) NF-κB-associated inflammatory and stress-responsive genes; (4) interferon signaling components; and (5) non-ISG transcriptional responses. At 6 hpi, cbVG⁺ cells expressed higher levels of these genes likely due interferon signaling originating from the cbVG⁺ subset. By 12 and 24 hpi, these differences diminished as shared antiviral programs spread across the population. To assess whether broad induction of these genes is cbVG-dependent, we performed qPCR validation and found that OASL, IFIT1, HERC5, and TNFAIP3 were upregulated only in cbVG-high infections, but not in the cbVG-clean stock (Fig. 6B-E), demonstrating that even population-wide antiviral programs require cbVGs for initial induction. We also identified genes that were not broadly upregulated at 6 hpi but became strongly induced in both cbVG⁺ and cbVG⁻ cells at 12 and 24 hpi (Fig. 6F). The functions of these genes are mainly related to interferon response and innate antiviral defense, cell surface signaling and immune interaction, transcriptional and stress-response regulation, vesicle trafficking and autophagy, metabolic and mitochondrial adaptation, and other cellular responses, suggesting that these genes contribute to a broader and more coordinated phase of host adaptation at later stages of infection. qPCR validation shown genes CXCL11, IFI27, and ITGA2 were upregulated in all cbVG-high stock infected cells but not in cells infected with the cbVG-clean stock or in mock controls (Fig. 6G-I).

**Fig. 6.**
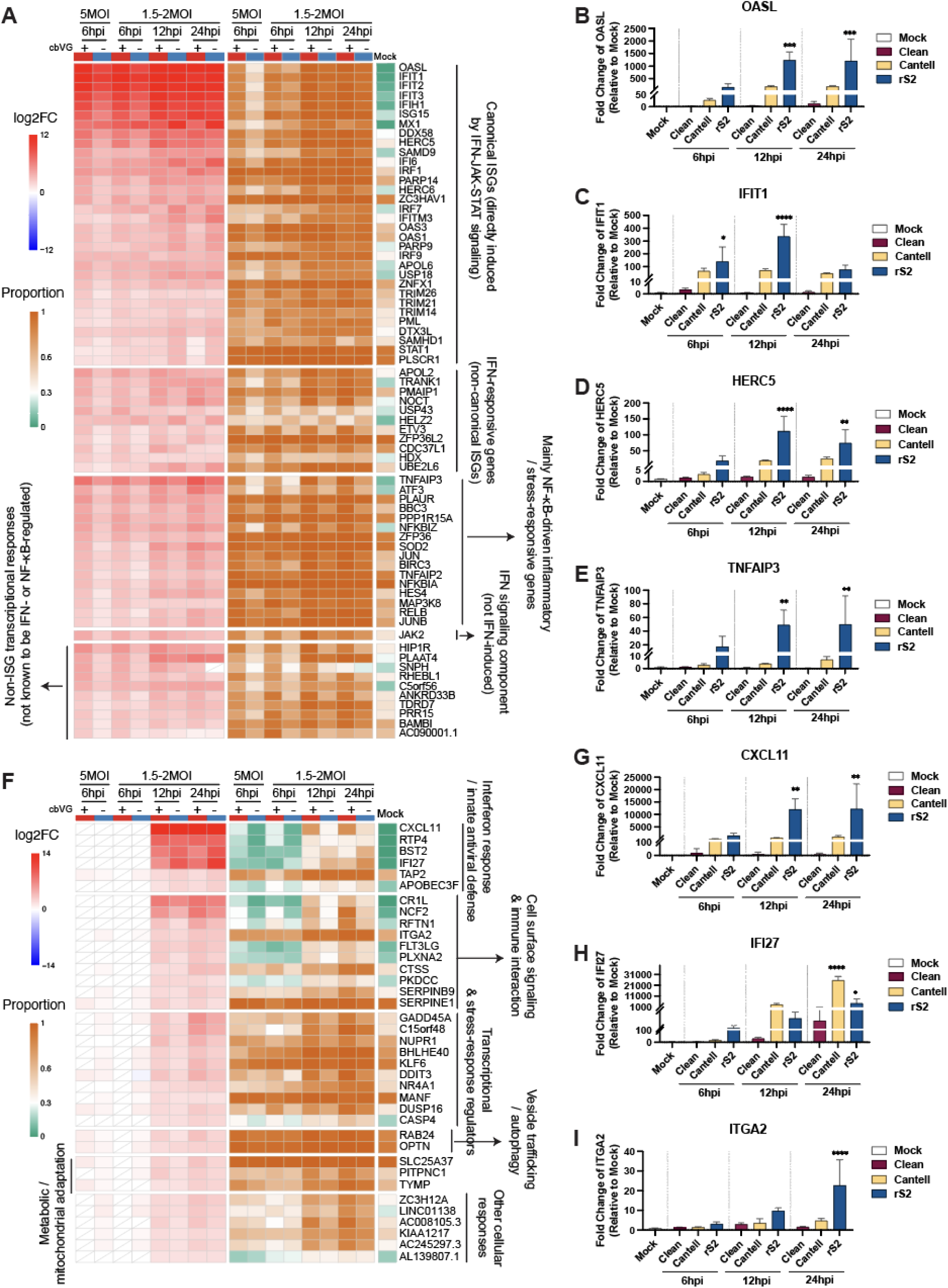
Functional classification and validation of genes upregulated in cbVG⁺ and cbVG⁻ cells. (A) Heatmap showing genes specifically upregulated in cbVG⁺ and cbVG⁻ cells at both 6hpi samples. Genes were grouped into five functional categories based on their transcriptional induction mechanisms. The red-blue color scale indicates the log2 fold change (log2FC), while the orange-green color scale represents the proportion of cells expressing each gene. White tiles with diagonal lines indicate features that did not meet the predefined adjusted P-value and proportion thresholds. (B-E) Representative genes validated by qPCR. A549 cells were infected with SeV Cantell and rS2. Each gene mRNA levels were quantified by qPCR and are presented as fold change relative to mock-infected samples. Cells were collected at 6, 12, and 24 hpi. Data represent mean ± SD from three biological replicates. Ordinary one-way ANOVA followed by Dunnett’s multiple comparisons test comparing each infection group with the mock control (n = 3). Adjusted P values are shown. Significance P values are indicated as follows: P < 0.05 (*), < 0.01 (**), < 0.001 (***), < 0.0001 (****). (F) Heatmap showing genes specifically upregulated in cbVG⁺ and cbVG− cells at both 12 hpi and 24 hpi, but not at 6 hpi. Genes were grouped into seven functional categories based on their known or predicted biological functions. The red-blue color scale indicates the log2 fold change (log2FC), while the orange-green color scale represents the proportion of cells expressing each gene. White tiles with diagonal lines indicate features that did not meet the predefined adjusted P-value and proportion thresholds. (G-I) Representative genes validated by qPCR. A549 cells were infected with SeV Cantell and rS2. Each gene mRNA levels were quantified by qPCR and are presented as fold change relative to mock-infected samples. Cells were collected at 6, 12, and 24 hpi. Data represent mean ± SD from three biological replicates. Ordinary one-way ANOVA followed by Dunnett’s multiple comparisons test comparing each infection group with the mock control (n = 3). Adjusted P values are shown. Significance P values are indicated as follows: P < 0.05 (*), < 0.01 (**), < 0.001 (***), < 0.0001 (****).

Together, these data define a shared antiviral program that becomes established across the infected population, but whose timing, magnitude, and breadth depend on early cbVG signals from a rare initiating subset of cells.

### cbVG⁺ cells undergo late-stage suppression of structural and metabolic programs

We then examined downregulated genes to assess whether cbVGs repress cellular functions during infection. At 6 hpi, no genes were consistently downregulated across cbVG⁺ cells, however 44 genes were significantly downregulated specifically in cbVG⁺ cells at 12 and 24 hpi (Fig. 7A). Some of these genes were also downregulated in the 6 hpi high-MOI condition, suggesting dose-dependent early repression. These repressed genes encompassed cell-surface signaling and adhesion, metabolic and mitochondrial adaptations, cytoskeletal organization, transcriptional and stress-response regulation, and vesicle trafficking, indicating that cbVG⁺ cells undergo structural and metabolic remodeling at late stages of infection. This shift may reflect transition from active antiviral defense toward cellular recovery mechanisms. We attempted to validate downregulated genes by qPCR, however the low levels of expression of these genes and the impact of the observed upregulation of some of the genes in cbVG⁻ cells make this validation approach impossible.

**Fig. 7.**
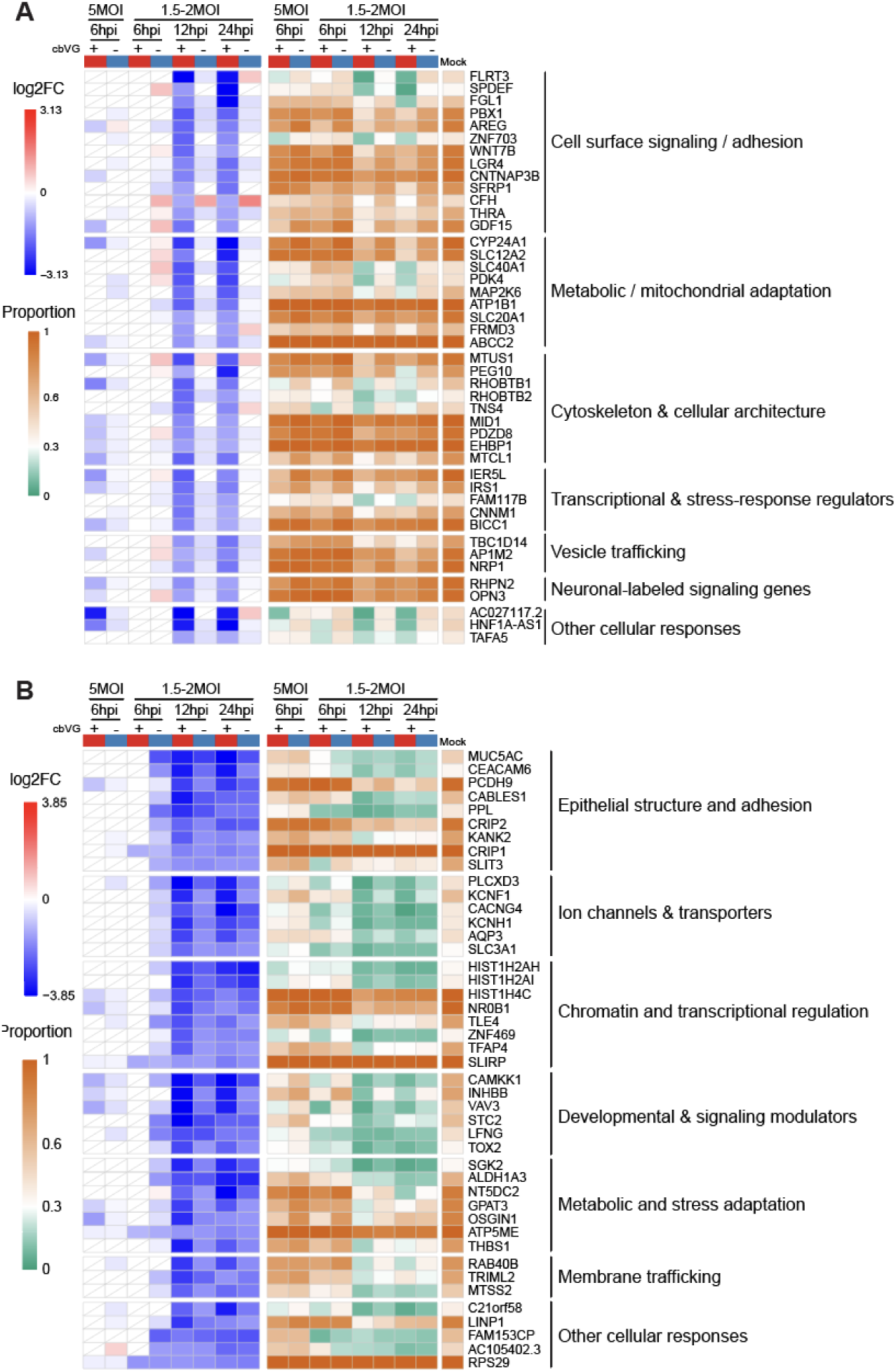
Functional classification and validation of genes downregulated at late infection. (A) Heatmap showing genes specifically downregulated in cbVG⁺ but not cbVG⁻ cells at both 12hpi and 24hpi samples. Genes were grouped into seven functional categories based on their known or predicted biological functions. The red-blue color scale indicates the log2 fold change (log2FC), while the orange-green color scale represents the proportion of cells expressing each gene. (B) Heatmap showing genes specifically downregulated in cbVG⁺ and cbVG⁻ cells at both 12hpi and 24hpi samples but not common at 6hpi. Genes were grouped into seven functional categories based on their known or predicted biological functions. The red-blue color scale indicates the log2 fold change (log2FC), while the orange-green color scale represents the proportion of cells expressing each gene. White tiles with diagonal lines indicate features that did not meet the predefined adjusted P-value and proportion thresholds.

**Fig. 8.**
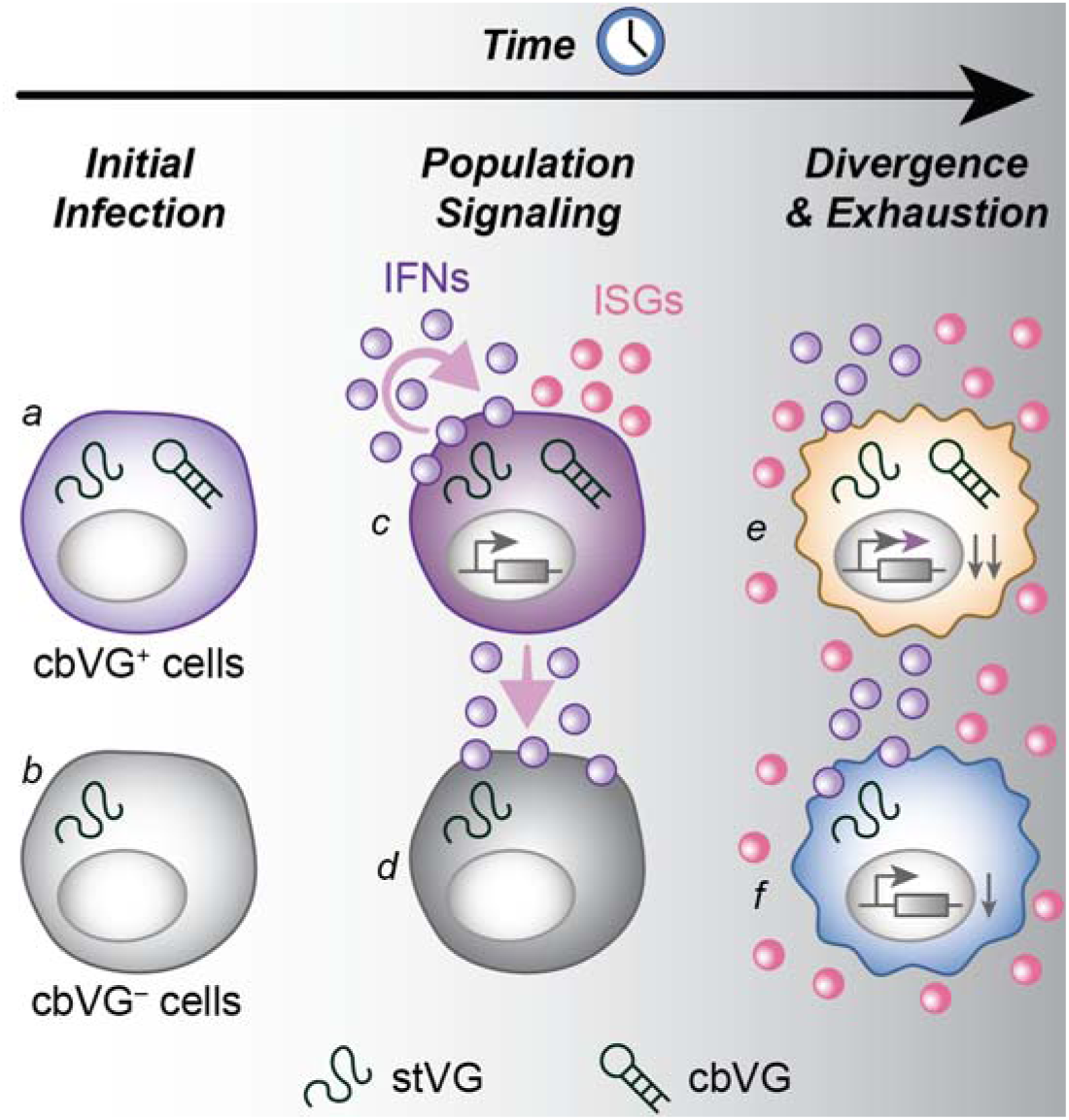
Model summarizing the role of cbVGs in driving antiviral responses during infection. At initial infection, there are two types of infected cells: cbVG⁺ cells (a) and cbVG⁻ cells (b). cbVG⁺ cells act as early initiators of innate immune signaling, inducing strong interferon (IFN) and interferon-stimulated gene (ISG) responses (c) and transmitting antiviral signals to neighboring cbVG⁻ cells through paracrine signaling (d). As infection progresses, IFN and ISG expression becomes broadly activated across the infected cell population (e, f). At later stages, cbVG⁺ cells additionally initiate unique transcriptional programs (e), while global transcriptional repression is observed in all infected cells and is stronger in cbVG⁺ cells (e) than in cbVG⁻ cells (f).

Across all SeV-infected groups, we identified another set of genes with reduced expression at 12 and 24 hpi relative to mock (Fig. 7B). Most of these genes showed stronger downregulation in cbVG⁺ cells, suggesting that cbVGs enhance late-stage suppression of cellular programs. These findings indicate that cbVG⁺ cells undergo late-phase structural and metabolic remodeling, reflecting a transition away from proliferative and epithelial programs during infection.

### A multilayered model of cbVG-driven host responses

Collectively, our single-cell analyses reveal a tightly regulated, multilayered architecture of host responses driven by cbVGs: Layer 1 (Origin layer, early): a rare cbVG⁺ population initiates interferon and chemokine programs, acting as the source of antiviral signaling within the infected population. Layer 2 (Amplified layer): these antiviral signals propagate to cbVG⁻ cells and establish a population-wide antiviral state, dominated by canonical ISGs and innate signaling programs. Layer 3 (Divergent layer, late): cbVG⁺ cells subsequently acquire distinct late-phase transcriptional programs related to cell-state remodeling, and stress adaptation. Layer 4 (Repression layer): both cbVG⁺ and cbVG⁻ cells undergo late-stage downregulation of structural, metabolic, and developmental programs, with stronger repression in cbVG⁺ cells. Thus, cbVGs determine not only the magnitude of antiviral responses, but also the timing, distribution, and differentiation of cell states during infection. These findings provide a mechanistic framework for understanding how cbVGs shape infection outcomes by orchestrating sequential transcriptional programs and guiding cell-state transitions within infected populations.

## Discussion

In this study, we developed a single-cell RNA sequencing workflow combined with a computational approach that enables direct detection of cbVGs and the host responses they induce at single-cell resolution. Because cbVG abundance is quantified using UMIs rather than total amplified reads, cbVG reads at single-cell resolution are less than in bulk RNA sequencing. This strategy reveals that cbVGs orchestrate the antiviral response and drive cell-state transitions during infection, providing the first high resolution view of cbVG-associated cellular response during a viral infection. Rather than uniformly activating antiviral programs, cbVGs initiate interferon and chemokine signaling in a rare subset of infected cells, which subsequently transmit these signals to surrounding cells and establish a second wave of antiviral transcriptional responses. This layered architecture indicates that cbVGs determine the timing, distribution, and magnitude of antiviral programs across the infected population.

Previous bulk measurements have shown that cbVGs can dominate antiviral signaling^9,13,35,58^, but these approaches mask cell-to-cell heterogeneity and obscure whether cbVGs simply correlate with antiviral activation or directly instruct population-level responses. At single-cell resolution, our analysis resolves how these population-level antiviral responses are organized, revealing that cbVG⁺ cells act as early initiators, whereas cbVG⁻ cells function as receivers that later transition toward inflammatory programs. Integrated clustering further revealed five transcriptional states whose relative abundance shifts over time. Using differential expression analyses of cbVG⁺ and cbVG⁻ cells, we identified gene sets that reflect distinct layers of cbVG-associated responses. Together, these single-cell-resolved findings provide a framework for interpreting the widespread presence of cbVGs in clinical infection across diverse RNA viruses^2,7,16,18,19,21,57,59^.

To date, however, most *in vivo* and clinical observations have established correlative relationships between cbVG abundance and disease outcome, viral loads, or antiviral responses^7,21,57,59^, without resolving how cbVGs shape host responses at the level of individual cells. In addition, cbVGs have been shown to spread within infected tissues^18^ and to associate with interferon and inflammatory signatures^7,21^, but whether these effects reflect direct sensing of cbVGs or secondary propagation of antiviral signals has remained unclear. Here, to directly define cbVG-associated host responses, we intentionally adopted a simplified experimental system using the human lung epithelial cell line A549. While this approach enabled us to isolate transcriptional programs tightly linked to cbVGs, it does not capture the full complexity of clinical infection. *In vivo*, virus infection including cbVG activity is likely influenced by host-specific variables such as age, sex, genetic background, and prior immune history^60–64^, as well as by the functional specialization of distinct cell types as they play different roles during virus infection^65,66^.

These considerations raise important unresolved questions regarding cbVG biology *in vivo*, including whether cbVGs preferentially arise or accumulate within specific cell types, and whether distinct cell populations exhibit differential sensitivity or transcriptional responses to cbVG-mediated signaling. Addressing these questions will require extending our findings along two complementary directions. First, the cbVG-associated gene sets identified here provide a resource for analyzing clinical samples, enabling evaluation of how these transcriptional programs correlate with cbVG abundance and clinical outcomes across patient cohorts. Second, *in vivo* infection models using cbVG-high viral stocks will be essential to determine whether cbVGs preferentially enrich within particular cell types and to directly compare cbVG⁺ and cbVG⁻ responses within defined cellular compartments. Notably, because sensitive and comprehensive detection of diverse cbVG species in complex tissues remains technically challenging, the gene signatures defined in this study may also serve as indirect readouts for cbVG activity. For example, cells co-expressing multiple genes we identified from only cbVG⁺ cells could be classified as functionally cbVG-positive, enabling identification of putative false-negative cells in which cbVGs are present but fall below current detection thresholds. Together, these approaches will help bridge single-cell mechanistic insights with the complexity of cbVG biology *in vivo* and in clinical infection. While our study focuses on epithelial cells, the conceptual separation of cbVG-initiating and cbVG-receiving states provides a framework that can be tested across additional cell types *in vivo*, including immune and stromal populations.

Our study also highlights technical considerations that will be important for future work. Mock controls were collected at 6 hpi, and epithelial gene expression can vary over time due to basal cycling, stress responses, or culture conditions, which may influence identification of late-regulated genes. Also, to minimize the effects of extensive cell death at later stages of infection, our analyses were limited to samples collected up to 24 hpi; therefore, potential cellular states emerging at later time points, such as the establishment of persistent infection in cbVG⁺ cells^35^, were not captured and should be considered in future studies. Additionally, standard scRNA-seq platforms capture only short sequences near the template-switch oligo, which limits detection of diverse cbVG species and can lead to false-negative classification of some cbVG⁺ cells. This is not an issue for SeV Cantell infection, which is dominated by a single abundant cbVG species (cbVG-546), but it becomes problematic for other virus stocks that produce a wider variety of cbVGs. Despite these limitations, the cbVG-associated gene signatures identified here provide a complementary framework for interpreting cbVG activity at the single-cell level, particularly in contexts where direct cbVG detection remains incomplete. As sequencing technologies continue to advance, long-read single-cell approaches may enable full-length capture of cbVGs and direct linkage of individual cbVG species to their induced transcriptional programs. Combining these technologies with cell-type-specific models will be essential for defining how cbVG generation, accumulation, and host responses differ across diverse tissues and for resolving the molecular basis of how cbVGs shape infection dynamics in complex biological settings.

## Materials & methods

### Cells and virus

A549 cells (ATCC, #CCL-185) were cultured in tissue culture medium (Dulbecco’s modified Eagle’s medium (DMEM) (Invitrogen, #11965092) supplemented with 10% fetal bovine serum (FBS) (Sigma, #F0926), gentamicin 50ug/ml (ThermoFisher, #15750060), L-glutamine 2 mM (Invitrogen, #G7513) and sodium pyruvate 1 mM (Invitrogen, #25-000-C1) at 5% CO2 37°C. Cells were treated with mycoplasma removal agent (MP Biomedical, #3050044) and tested monthly for mycoplasma contamination using the MycoAlert Plus mycoplasma testing kit (Lonza, #LT07-318). SeV Cantell strain was propagated in 10-day-old embryonated chicken eggs (Charles River Laboratories) for 40 hours, as previously described ^67^. SeV clean stocks and cbVG-high stock rS2 were rescued from cDNA using reverse genetics and amplified in LLCMK2 cells. Their cbVG abundance and species composition were characterized as previously described ^14^.

### Virus infection and samples preparation for scRNA-Seq

Infections were performed after washing the cells once with PBS and incubating them with virus diluted in infection media (DMEM, 35% bovine serum albumin (Sigma, #A7979), penicillin-streptomycin (Gibco, #15140-122), and 5% NaHCO₃ (Gibco, #25080094)) at 37°C for 1 hour, with gentle shaking every 15 minutes. Cells were then washed twice with PBS and supplemented with fresh infection media. The infected cells were incubated at 37°C and harvested at 6, 12, and 24 hpi. Mock-infected cells (infection media only) were used as controls. For each infection, two replicates were included. Infected or mock-infected cells were collected by detachment with Trypsin-EDTA 0.05% (Thermo, # 25300054) and subjected to live/dead staining (Thermo Fisher, # 65-0866-14), followed by flow cytometry sorting to isolate live cells.

### Library preparation and sequencing

Live cells were resuspended in 1% BSA in 1× PBS at a concentration of 1,000 cells/µL, and 15 µL from each sample was used for library preparation. For each sample, 35.2 µL of a virus-specific RT primer (10 uM; 5′ - AAGCAGTGGTATCAACGCAGAGTACTGCTCCTCAGGGTGGATACT-3 ′ ) was added in combination with the conventionally used oligo-dT primer at an equal volume and concentration. Thus, both the virus-specific RT primer and oligo-dT were included during reverse transcription to simultaneously capture viral genomes/cbVGs and poly-adenylated host and viral transcripts. Library preparation was performed following the 10x Genomics Chromium Next GEM Single Cell 5′ v2 protocol. Samples were sequenced on an Illumina NovaSeq 6000 to generate 150 bp paired-end reads, targeting 700 million reads per sample, at the Genome Access Technology Center (GTAC) core sequencing facility at Washington University. Raw sequencing data have been deposited in the NCBI Sequence Read Archive (SRA) under BioProject accession number PRJNA1390230.

### Upstream computational analyses of scRNA-Seq data

Raw Illumina sequencing data were aligned to the human reference genome (GRCh38) using Cell Ranger (v7.2.0) to generate gene count matrices, which were used to create Seurat objects for host gene expression analysis. Reads that did not map to the human genome were extracted from the Cell Ranger output and aligned to the SeV genome using Cell Ranger with a custom-built reference. In parallel, cbVG junction reads were identified using VODKA2 following specific preprocessing steps^30^. Briefly, the template switching oligo (TSO) sequence was removed from R1 using fastx_trimmer, and Illumina adapters, oligo(dT) tails, and 5′ overhang sequences were trimmed from both R1 and R2 using cutadapt. The resulting SeV standard genome and cbVG read counts per barcode were then added to the Seurat objects as metadata. Doublet cells were identified and removed using the scDblFinder R package, and low-quality cells were filtered based on quality control (QC) metrics using a sample specific approach. Filtering thresholds were defined as the median ± 3 × median absolute deviation (MAD) for key QC metrics, utilizing the outlier detection logic implemented in the isOutlier() function from the scater package^68^. Specifically, cells were excluded if their mitochondrial transcript proportion was upward outlier, or if their total RNA counts or number of detected features were downward outliers. All downstream analyses and visualization were performed in R (v4.4 or 4.5).

### Clustering in Seurat

Per-sample clustering: Each sample was processed independently to capture sample-specific heterogeneity. After normalization and variance stabilization using SCTransform(), principal component analysis (PCA) was conducted, followed by dimensionality reduction via Uniform Manifold Approximation and Projection (UMAP). Clustering was performed using the Seurat FindClusters() function at a resolution of 0.2 or 0.4. Differential gene expression (DGE) analysis was performed to characterize distinct cell populations within each sample. Integrated clustering: To identify shared transcriptional programs across timepoints, we performed integrated clustering on specific cell subsets (e.g. all SeV-positive cells or across all SeV-negative cells) from infected samples. Datasets were integrated using the standard Seurat anchor-based strategy. Following SCTransform on individual samples, we identified common integration features across datasets, ran PrepSCTIntegration() and identified anchors using FindIntegrationAnchors(). After merging the data with IntegrateData(), the integrated object was then subjected to PCA and UMAP reduction. Shared cell states were identified using a clustering resolution of 0.2 and DGE analysis was conducted across these integrated clusters to identify conserved transcriptional modules.

### Gene expression analysis of cbVG⁺ and cbVG⁻ cells relative to mock

To identify cbVG-specific transcriptional programs, SeV-infected cells were stratified into cbVG⁺ and cbVG⁻ populations based on the detection of cbVG junction sequences. DGE analysis was performed using Seurat FindMarkers() to compare cbVG⁺ versus mock and cbVG⁻ versus mock uninfected cells independently. Due to technical limitations in cbVG detection, a small number of cbVG⁺ cells may be included in the cbVG⁻ population; therefore, we applied a percentage threshold for DGE analysis. Genes were defined as significantly upregulated in cbVG⁺ or cbVG⁻ cells if they met the following criteria: detected in >30% of cells in the population (min.pct > 0.3) and adjusted p-value < 0.01, and log2 fold-change > 1. Conversely, genes were defined as significantly downregulated relative to mock if they were detected in > 30% of mock cells, had an adjusted p-value < 0.01, and log2 fold-change < -1. This analytical design distinguishes cbVG-dependent responses from changes driven by SeV infection alone.

### Genes validation by RT-qPCR

A549 cells were infected with SeV stocks (clean stock, Cantell, and rS2) at a defined MOI and harvested at 6, 12, and 24 hpi. Each infection condition included three biological replicates. Total RNA from infected and control samples was extracted using a KingFisher instrument and the MagMAX mirVana Total RNA Isolation Kit (Thermo Fisher, #A27828) following the manufacturer’s guidelines. A total of 150 ng RNA was used for cDNA synthesis with the High-Capacity RNA-to-cDNA Kit (Thermo Fisher, #18080051). qPCR was performed using SYBR Green (Thermo Fisher, # A46113) and 5 µM of forward and reverse primers for each gene on an Applied Biosystems QuantStudio 5 machine. Primers used for qPCR are listed in Supplemental Table 3. Relative gene expression was normalized to human GAPDH and human beta-actin expression as described previously^69^. qPCR reactions were performed in technical triplicates and data represent the mean of biological replicates.

## Supporting information

All supplementary figures

## Acknowledgments

We thank the Genome Technology Access Center (GTAC) and Jennifer Ponce at the McDonnell Genome Institute at Washington University School of Medicine in St. Louis for assistance with library preparation and sequencing. We thank Dr. Jan Carette and Dr. Allison Dupzyk from the Department of Microbiology and Immunology at Stanford University for their early technical advice on virus-specific primer design. We also thank the Siteman Cancer Center Flow Cytometry Core at Washington University School of Medicine in St. Louis for assistance with sorting infected cells. This work was funded by NIH/NIAID grants AI37062 and AI188900 and by a BJC Investigator Award to CBL. This work was also supported by the Alexander & Gertrude Berg Fellowship awarded to YY and Helen Hay Whitney Foundation Postdoctoral Fellowship awarded to MT.

## Declaration of generative AI and AI-assisted technologies in the manuscript preparation process

During the preparation of this work the authors used ChatGPT in order to improve the clarity and readability of the manuscript text. After using this tool, the authors extensively reviewed and edited the content as needed and take full responsibility for the content of the published article.

