## Supplementary material for "Detailed single-cell mapping of the transcriptional response to a virus infection driven by copy-back viral genomes": All supplementary figures

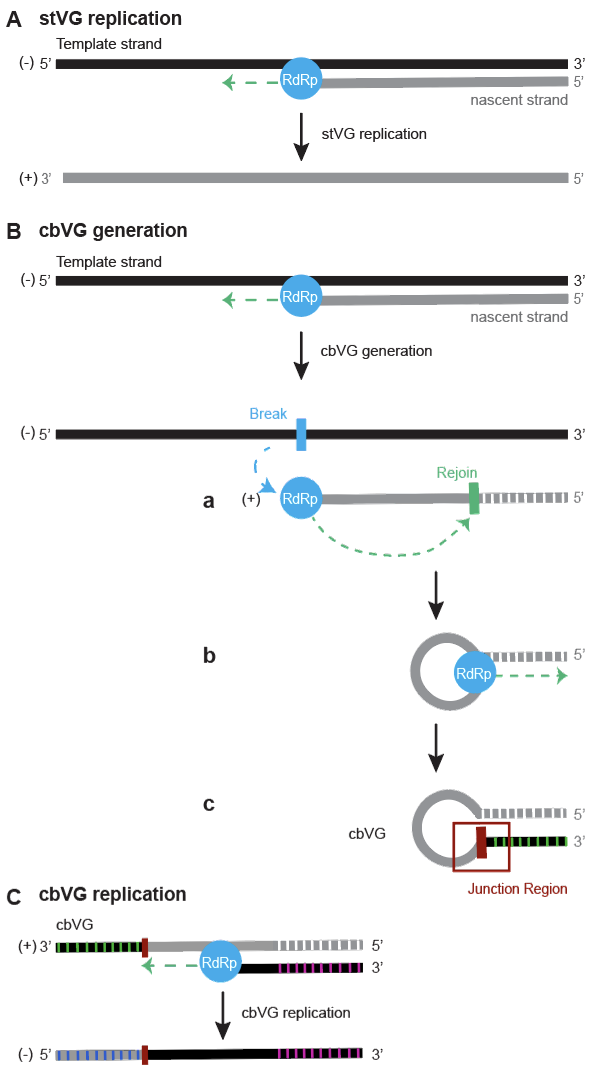

**Fig. S1. Visualization of the generation and replication of standard viral genomes (stVG) and copyback viral genomes (cbVG).** (A) stVG replication. The viral RNA-dependent RNA polymerase (RdRp, blue) synthesizes a full-length stVG using the negative-sense genomic RNA as a template. (B) cbVG generation. During replication, RdRp detaches from the template at a break point (blue) and rejoins the nascent strand at a rejoin position (green), forming a cbVG with complementary ends and a unique junction region (red). White stripes mark the template sequence used after the rejoin point, and green stripes indicate the complementary sequence at the 3′ end of the newly formed cbVG. (C) cbVG replication. cbVGs are replicated by RdRp. Pink and blue stripes indicate sequences complementary to the 3′ and 5′ ends of the cbVG template, respectively.

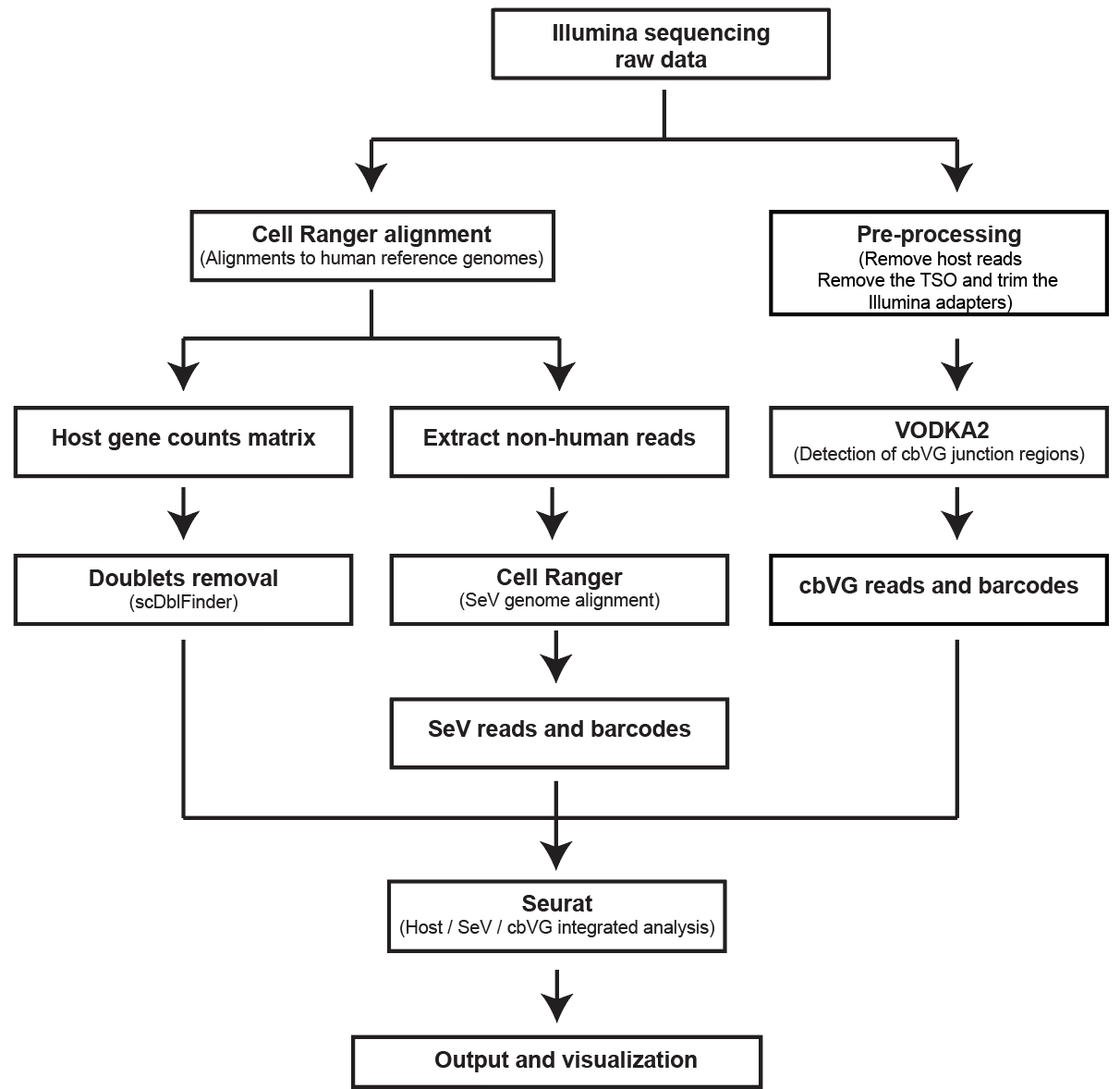

**Fig. S2. Workflow for detecting and integrating cbVGs from single-cell RNA-seq data.** Raw Illumina sequencing data were processed through parallel bioinformatic workflows for host and viral analyses. On the left, reads were aligned to the human reference genome using Cell Ranger, followed by generation of the host gene count matrix and doublet removal (scDblFinder). Non-human reads were extracted and re-aligned to the SeV genome to obtain SeV reads and barcodes. On the right, reads were pre-processed to remove host sequences, TSO, and Illumina adapters, then analyzed with VODKA2 for cbVG junction detection to obtain cbVG reads and barcodes. Host, SeV, and cbVG read count matrices were integrated by cell barcode using Seurat for multimodal analysis and visualization.

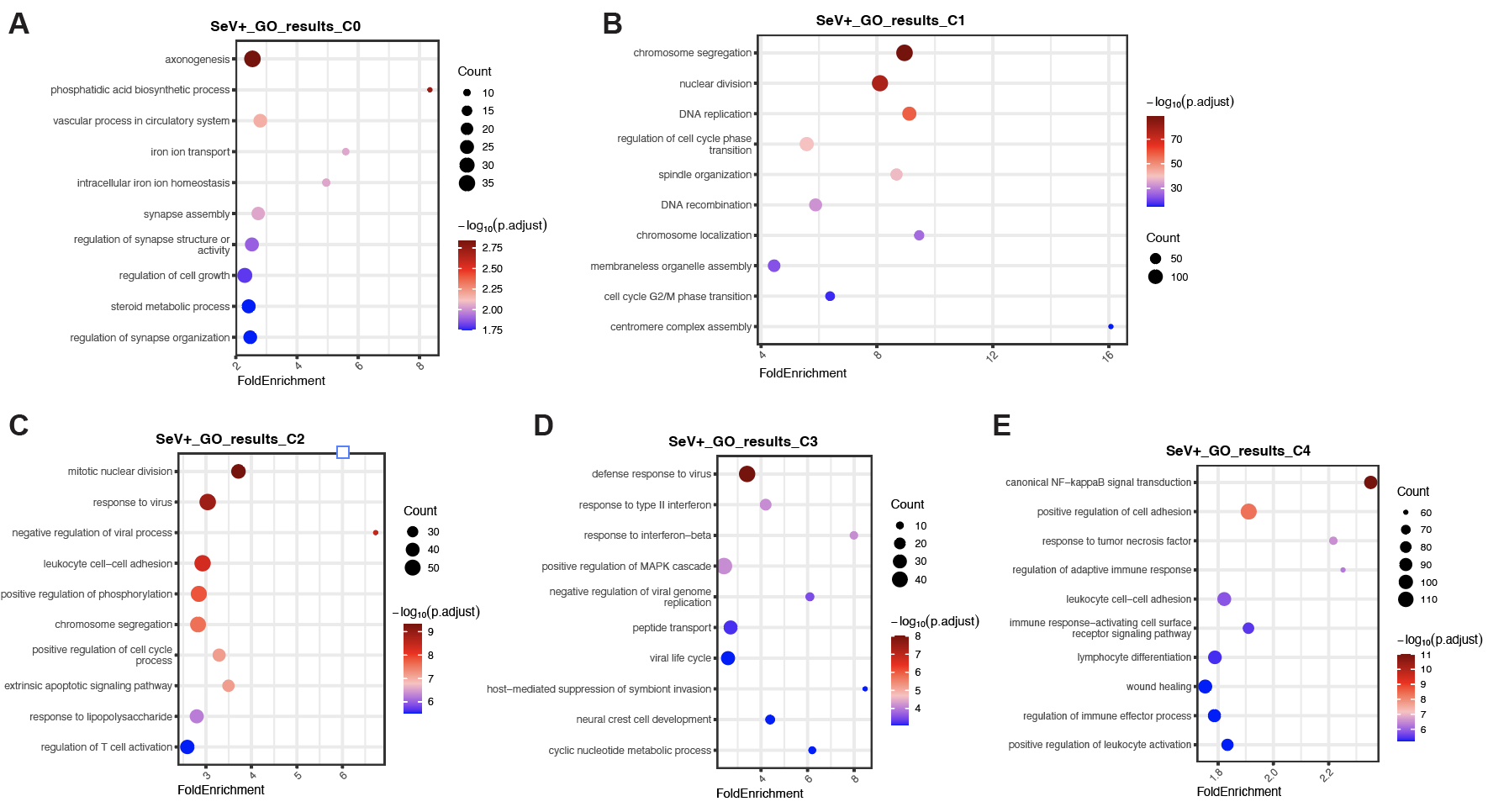

**Fig. S3. Gene Ontology (GO) enrichment analysis of SeV+ cell clusters.** Bubble plots showing the top enriched GO biological processes for marker genes of each SeV+ cluster (C0–C4). The x-axis indicates fold enrichment, the y-axis lists enriched GO terms, dot size represents the number of genes associated with each term, and color indicates the adjusted p-value (–log₁₀ p). (A) C0, (B) C1, (C) C2, (D) C3, and (E) C4.

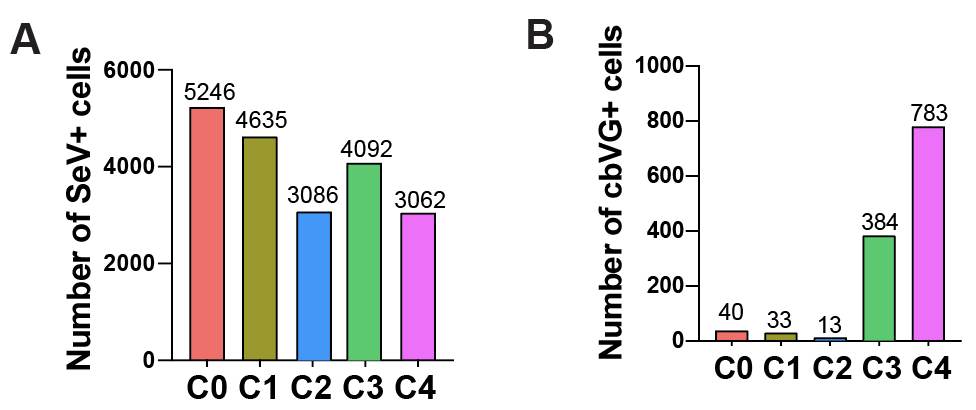

**Fig. S4. Number of cells in interegted analysis.** (A) Number of SeV+ cells in each cluster. (B) Number of cbVG⁺ cells in each cluster.

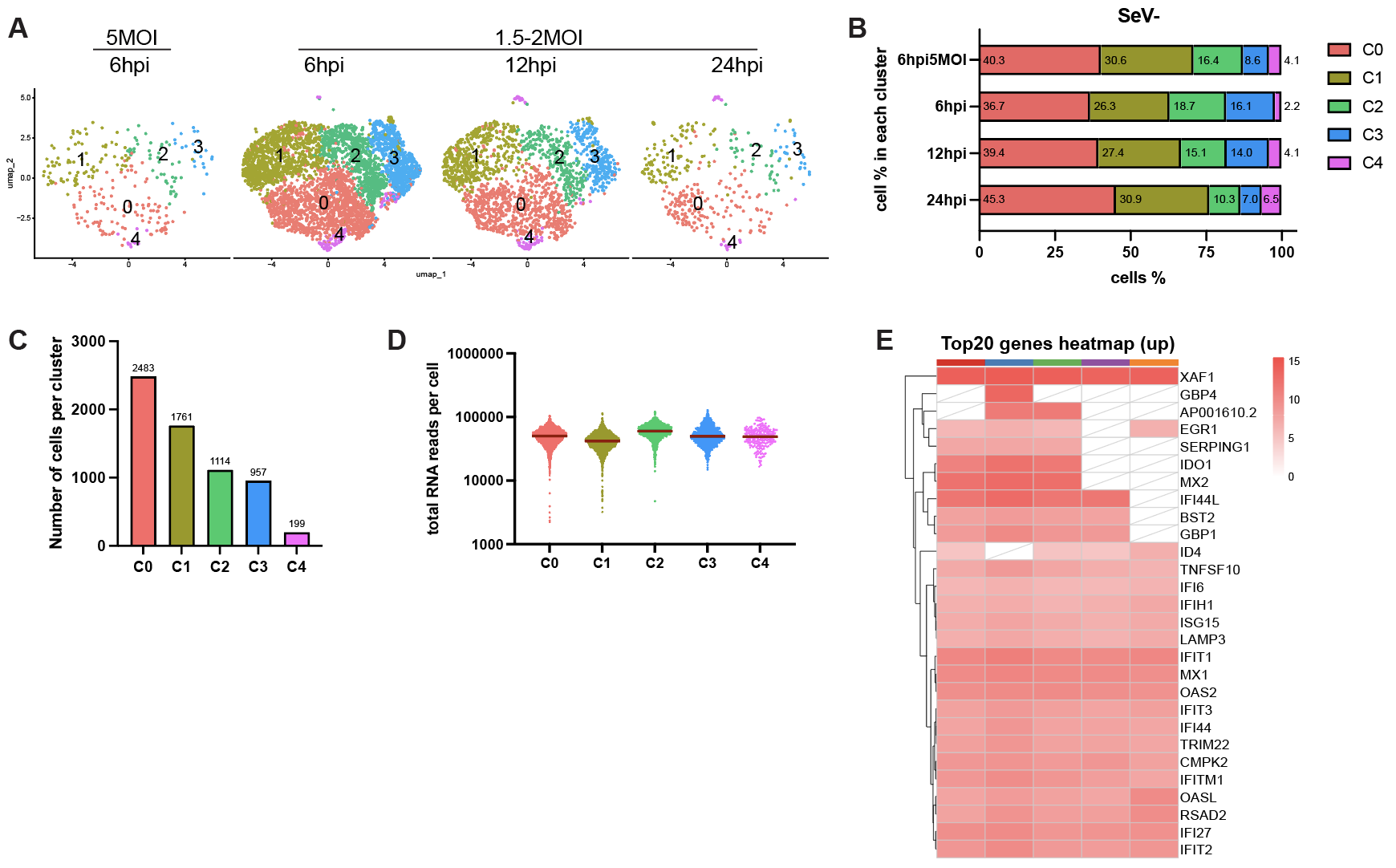

**Fig. S5. SeV- cells were divided into five populations based on host gene transcription.** (A) Integrated analysis of all SeV- cells from each sample. (B) Percentage of cells in each population across samples. (C) Number of SeV- in each cluster. (D) Total UMI counts per cell. The red line indicates the median. (E) Heatmap of the top 20 most upregulated genes from each cluster compared with mock. Average log2 fold change is shown as the scale bar. White tiles with diagonal lines indicate features that did not meet the predefined adjusted P-value and proportion thresholds.

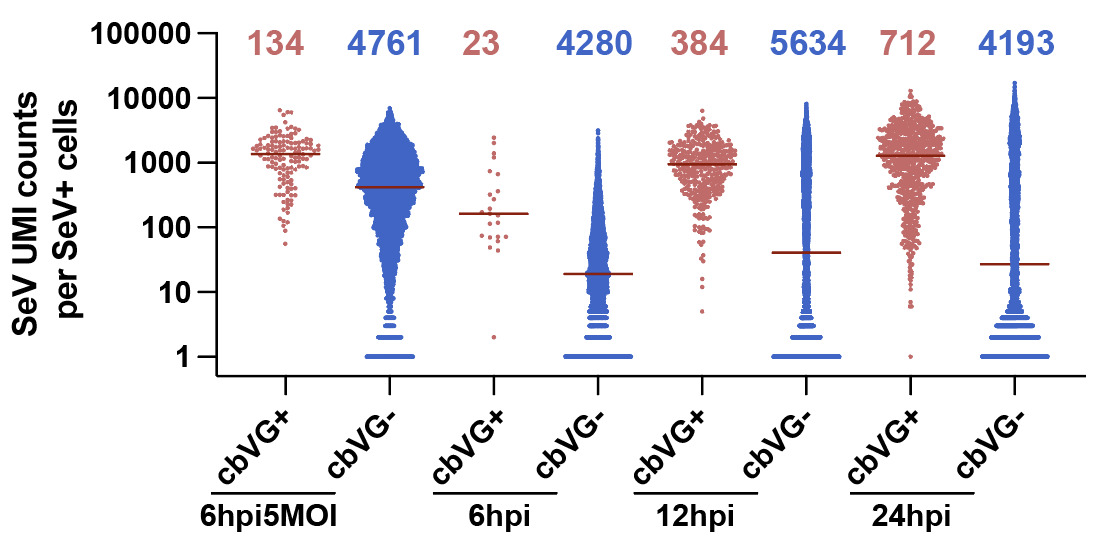

**Fig. S6.** **SeV UMI counts per cell in cbVG⁺ and cbVG⁻ groups.** Scatter plots showing SeV UMI counts per cell in cbVG⁺ (red) and cbVG⁻ (blue) populations at different condition. Each dot represents a single cell, and horizontal bars indicate median values. Numbers above each group denote the number of cells analyzed.

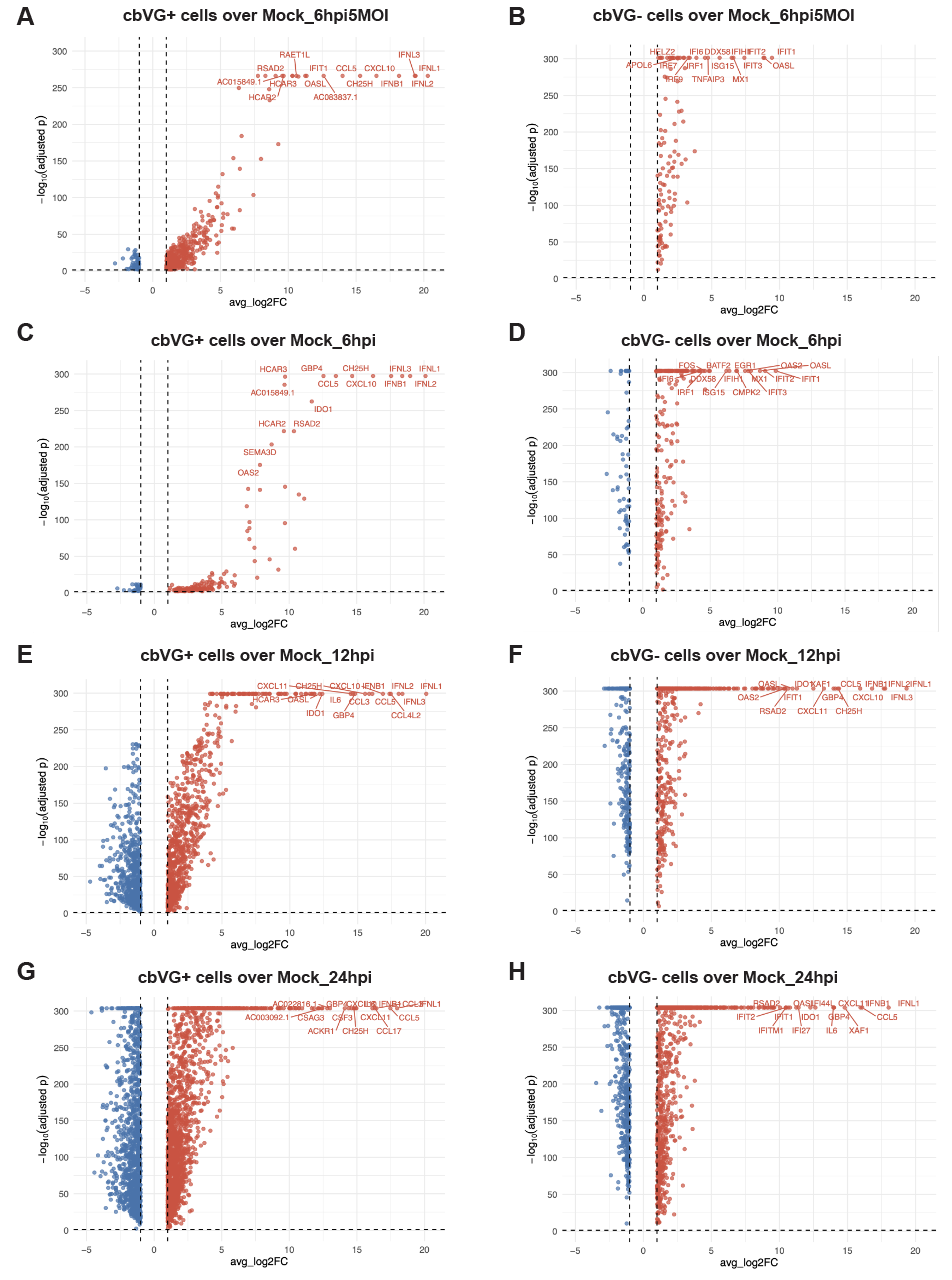

**Fig. S7.** **Differential gene expression in cbVG⁺ and cbVG− cells compared with mock (uninfected) cells.** Volcano plots showing differentially expressed genes between cbVG⁺ or cbVG^−^ cells and mock cells at the indicated infection stages: (A, B) 6 hpi5MOI, (C, D) 6 hpi, (E, F) 12 hpi, and (G, H) 24 hpi. Each dot represents one gene, with the x-axis showing average log2 fold change (avg_log2FC) and the y-axis showing -log10 adjusted p-value. Red dots indicate upregulated genes, and blue dots indicate downregulated genes in cbVG⁺ (left) or cbVG^−^ (right) SeV+ cells. For visualization, p values equal to 0 were replaced with one-tenth of the smallest non-zero p value across all genes.

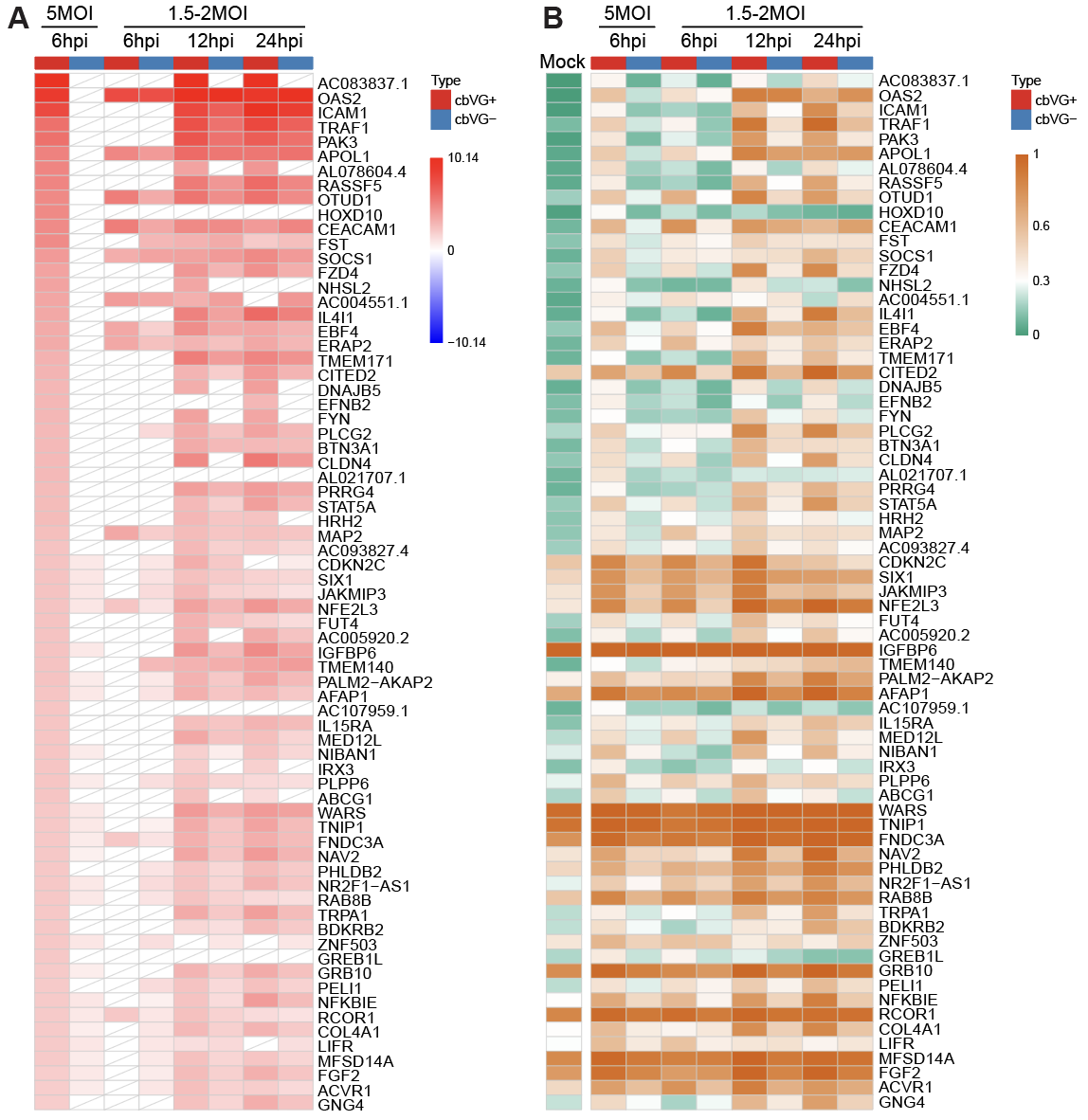

**Fig. S8. Expression levels and proportions of selected genes.** Heatmap showing genes specifically upregulated (lfc2>2) in cbVG⁺ 6hpi 5MOI samples. The red-blue color scale indicates the log2 fold change (log2FC) (A), while the orange-green color scale represents the proportion of cells expressing each gene (B). White tiles with diagonal lines indicate features that did not meet the predefined adjusted P-value and proportion thresholds.

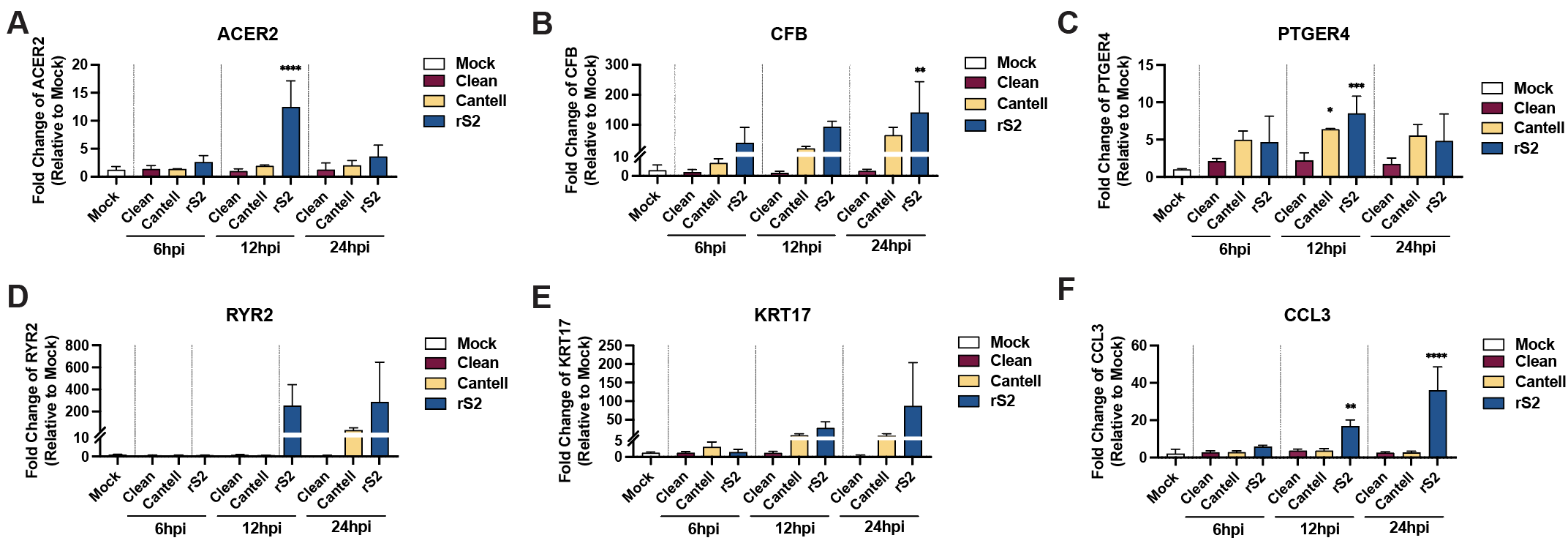

**Fig. S9. Representative genes validated by qPCR.** A549 cells were infected with SeV stocks containing different cbVG compositions: cbVG clean stock, Cantell, and rS2. Each gene mRNA levels were quantified by qPCR and are presented as fold change relative to mock-infected samples. Cells were collected at 6, 12, and 24 hpi. Data represent mean ± SD from three biological replicates. Ordinary one-way ANOVA followed by Dunnett’s multiple comparisons test comparing each infection group with the mock control (n = 3). Adjusted P values are shown. Significance P values are indicated as follows: P < 0.05 (*), < 0.01 (**), < 0.001 (***), < 0.0001 (****).

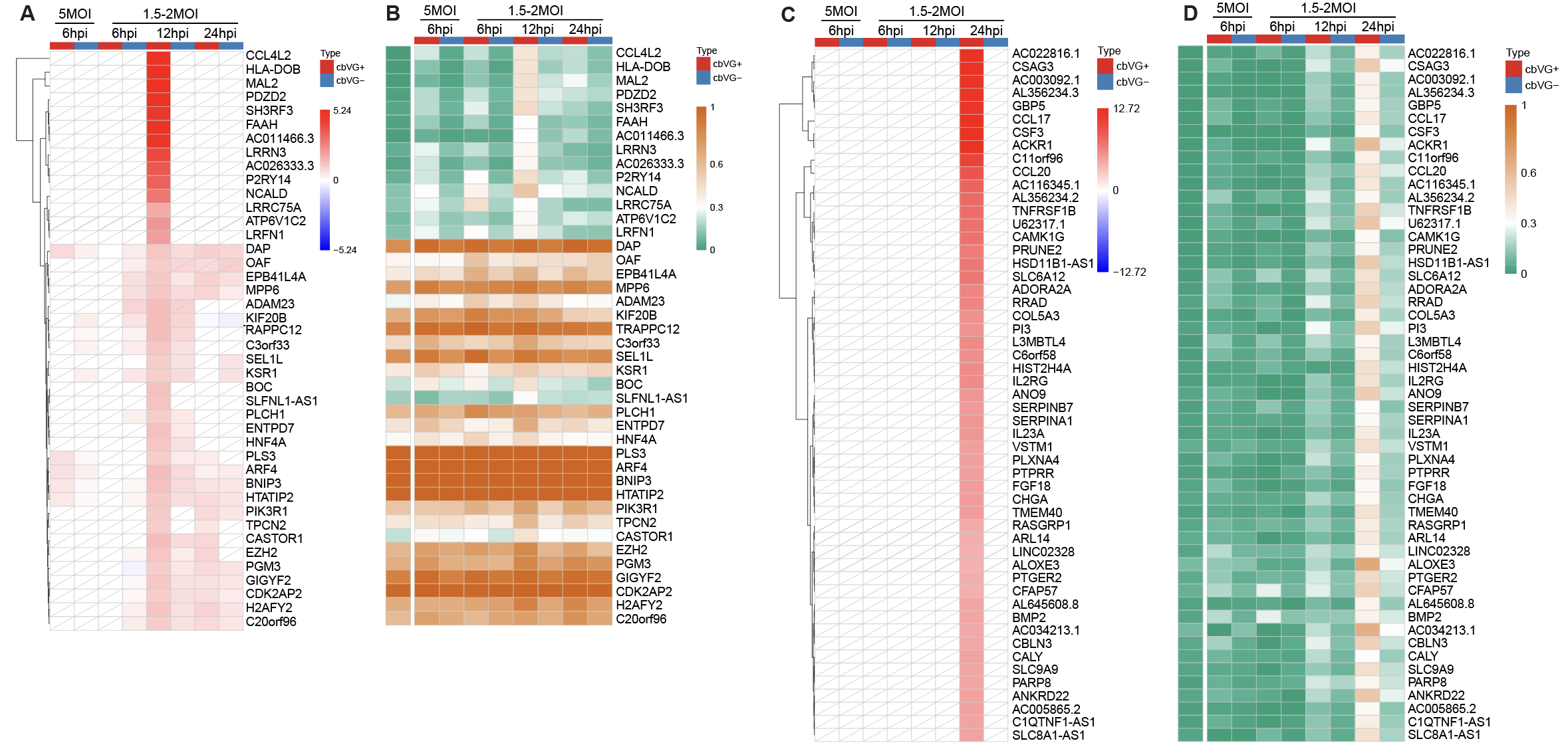

**Fig. S10. Expression levels and proportions of selected genes.** (A-B) Heatmap showing genes specifically upregulated in cbVG⁺ 12hpi samples. The red-blue color scale indicates the log2 fold change (log2FC), while the orange-green color scale represents the proportion of cells expressing each gene. (C-D) Heatmap showing genes specifically upregulated in cbVG⁺ 24hpi samples. The red-blue color scale indicates the log2 fold change (log2FC), while the orange-green color scale represents the proportion of cells expressing each gene. White tiles with diagonal lines indicate features that did not meet the predefined adjusted P-value and proportion thresholds.
